# Structures of the substrate-engaged 26S proteasome reveal the mechanisms for ATP hydrolysis-driven translocation

**DOI:** 10.1101/393223

**Authors:** Andres H. de la Peña, Ellen A. Goodall, Stephanie N. Gates, Gabriel C. Lander, Andreas Martin

## Abstract

The 26S proteasome is the primary eukaryotic degradation machine and thus critically involved in numerous cellular processes. The hetero-hexameric ATPase motor of the proteasome unfolds and translocates targeted protein substrates into the open gate of a proteolytic core, while a proteasomal deubiquitinase concomitantly removes substrate-attached ubiquitin chains. However, the mechanisms by which ATP hydrolysis drives the conformational changes responsible for these processes have remained elusive. Here we present the cryo-EM structures of four distinct conformational states of the actively ATP-hydrolyzing, substrate-engaged 26S proteasome. These structures reveal how mechanical substrate translocation accelerates deubiquitination, and how ATP-binding, hydrolysis, and phosphate-release events are coordinated within the AAA+ motor to induce conformational changes and propel the substrate through the central pore.

---

The 26S proteasome is the final component of the ubiquitin-proteasome system and thus central to general proteostasis and the regulation of numerous vital processes in eukaryotic cells^1^. Proteins are targeted for proteasomal degradation through the covalent attachment of poly-ubiquitin chains to lysine residues^2^. To safeguard against indiscriminate degradation, the proteolytic active sites of the proteasome are sequestered within the barrel-shaped 20S core particle (CP). Access to these active sites is controlled by the 19S regulatory particle (RP), which binds to one or both ends of the CP, recruits ubiquitinated proteins, and catalyzes their deubiquitination, unfolding, and translocation through a central pore into the proteolytic chamber of the CP for degradation^3^. The RP can be further subdivided into the base and lid subcomplexes. The 9-subunit lid subcomplex fulfills important scaffolding functions and contains the Zn^2+^-dependent deubiquitinase Rpn11, which is positioned above the central pore of the proteasome to remove ubiquitin chains from substrates prior to degradation^4-8^. The base subcomplex consists of ten subunits, including three ubiquitin receptors and six distinct AAA+ (ATPases Associated with diverse cellular Activities) ATPases, Rpt1-Rpt6^3,9^. These ATPases form a heterohexameric ring (in the order Rpt1, Rpt2, Rpt6, Rpt3, Rpt4, Rpt5) that represents the molecular motor of the proteasome^10^. Each Rpt consists of a N-terminal helix, an OB (oligonucleotide binding)-fold domain, and a C-terminal AAA+ motor domain. In the heterohexamer, the N-terminal helices of neighboring Rpt pairs form a coiled coil, and the six OB-fold domains assemble into a rigid N-ring above the AAA+ motor ring^6,8^. After ubiquitin-mediated substrate recruitment, the ATPase motor engages a flexible initiation region of the substrate for subsequent mechanical translocation and unfolding^11^. To facilitate substrate transfer to the CP, the ATPase hexamer also triggers opening of the CP access gate by docking conserved C-terminal tails of Rpt subunits into pockets at the surface of the CP α-ring^12-14^.

Like other AAA+ ATPases, the Rpt subunits contain a highly-conserved nucleotide binding pocket that couples ATP binding and hydrolysis with conformational changes to produce mechanical work^15,16^. This pocket is largely formed by the signature Walker-A and Walker-B motifs, responsible for nucleotide binding and hydrolysis, respectively, and an arginine finger provided by the clockwise-neighboring ATPase subunit that coordinates the γ-phosphate of ATP during hydrolysis and enables subunit communication^17^. Conserved pore-1 loops protruding from each ATPase subunit into the central channel sterically interact with substrate polypeptide and transduce nucleotide-dependent conformational changes into directional translocation^18-21^.

The common functional architecture of ring-shaped hexameric helicases and AAA+ translocases gave rise to a “ handover-hand” model for substrate translocation^22-24^, which is supported by numerous cryo-EM structures of substrate-bound homohexameric AAA+ motors^25-30^. Notably, these prior structures were trapped using hydrolysis-inactivating Walker-B mutations, non-hydrolyzable ATP analogs or analogs that are slowly hydrolyzed, to reveal series of subunits in the hexamer that resemble the ATP-bound, ADP-bound, and nucleotide-free states. Generally, five nucleotide-bound subunits contact the substrate polypeptide in a spiral-staircase arrangement of pore loops, whereas one subunit remains disengaged and nucleotide-free or ADP-bound. The “ hand-overhand” model stems from inferences regarding how individual subunits may progress through the various nucleotide states and substrate-binding conformations around the ring. The heterohexameric proteasomal AAA+ motor has been shown to adopt distinct spiral-staircase arrangements with individual Rpts in different vertical positions^6,31-35^, thus promising more detailed insights into the progression of states during the ATP-hydrolysis and substrate-translocation cycles. However, high-resolution structural studies of the proteasome during active substrate translocation, aimed at revealing numerous specific states of the dynamic ATPase motor, have so far been unsuccessful.

In the absence of substrate, the ATP-hydrolyzing proteasome primarily adopts the “ s1” state^6,36^ in which the ATPase domains of Rpt1-Rpt6 form a spiral staircase that is not coaxially aligned with the CP, and Rpn11 is positioned offset from the central pore of the motor. A low-resolution structure of the proteasome trapped with a stalled protein in the central pore revealed that upon substrate engagement, the RP transitions from the s1 state to a processing conformation, which is characterized by a more planar ATPase ring, a rotated lid subcomplex, and a coaxial alignment of Rpn11, the Rpt hexamer, and the CP^31^. However, the limited resolution and strong heterogeneity of the ATPases within these stalled proteasome complexes prevented the visualization of substrate and the identification of distinct motor states.

States that share structural similarities with the substrate-processing conformation are also observed for the substrate-free proteasome as a small sub-population in the presence of ATP (“ s2” state) and upon ATPase inhibition using either ATP analogs or Walker-B mutations in individual Rpt subunits (“ s3”, “ s4”, “ s5”, “ s6” states; and the unnamed state seen in ADP-AlF_x_)^14,33-35^. Cryo-EM reconstructions of these states revealed distinct spiral-staircase arrangements and nucleotide occupancies of Rpt subunits, but the lack of ATP hydrolysis and the absence of substrate limited the conclusions that could be drawn regarding the mechanisms for ATP-hydrolysis-coupled translocation.

We explored the mechanistic details of ATP-hydrolysis-driven substrate translocation by determining the structure of the substrate-engaged 26S proteasome in the presence of ATP. Unlike previous studies that utilized ATPase inhibition to trap substrate-bound states of other AAA+ motors, we stalled substrate translocation in the actively hydrolyzing motor of the proteasome by inhibiting Rpn11-mediated deubiquitination. We describe four cryo-EM structures, depicting four unique motor states with the unambiguous assignment of substrate polypeptide traversing the RP from the lysine-attached ubiquitin at the Rpn11 active site, through the Rpt hexamer, to the gate of the CP. Three of these states represent sequential stages of ATP binding, hydrolysis, and substrate translocation, and hence reveal, for the first time, the coordination of individual steps in the ATPase cycle of the AAA+ hexamer and their mechano-chemical coupling with translocation.

## Four substrate-bound 26S proteasome structures

To stall translocation at a defined substrate position, we inactivated the Rpn11 deubiquitinase of *Saccharomyces cerevisiae* 26S proteasomes by incubation with the inhibitor ortho-phenanthroline^4^ and added a globular model substrate with a single poly-ubiquitinated lysine flanking an unstructured C-terminal initiation region. Proteasomes thus engaged the flexible initiation region and translocated the substrate until the attached ubiquitin chain reached the inhibited Rpn11, preventing further translocation and trapping the substrate in the central pore, which is indicted by a complete inhibition of degradation (SFig. 1A). Importantly, stalling substrate translocation in the proteasome does not also stall the AAA+ motor, as we observed a rate of ATP hydrolysis that was even slightly elevated compared to freely translocating proteasomes (SFig. 1B). We posit that this stalled state resembles the scenario when the proteasome encounters thermodynamically stable substrate domains that require repeated pulling by the ATPase to be unfolded^31,37^.

Following incubation with substrate, proteasomes were vitrified for cryo-EM single particle analysis, which produced reconstructions of the 26S proteasome in six unique conformational states. In the initial 3D classification, roughly 42% of particles were observed to be substrate-free, adopting an s1-like state (SFig. 2A, Supplementary Table 1), while the rest of the particles were sorted into reconstructions that showed ubiquitin density adjacent to Rpn11, and accordingly adopted non s1-like conformations (SFig. 2A). Further focused classification of the ATPase motor resulted in four s4-like reconstructions and one reconstruction that resembled the s2 state, but lacked density for substrate within the central channel of the AAA+ motor. In contrast, the four s4-like reconstructions (ranging in overall resolution from ~4.2 to ~4.7 Å) showed clearly visible substrate density threaded through the center of the RP (Fig. 1, SFigs. 2, 3A and Supplementary Table 1).

**Figure 1.**
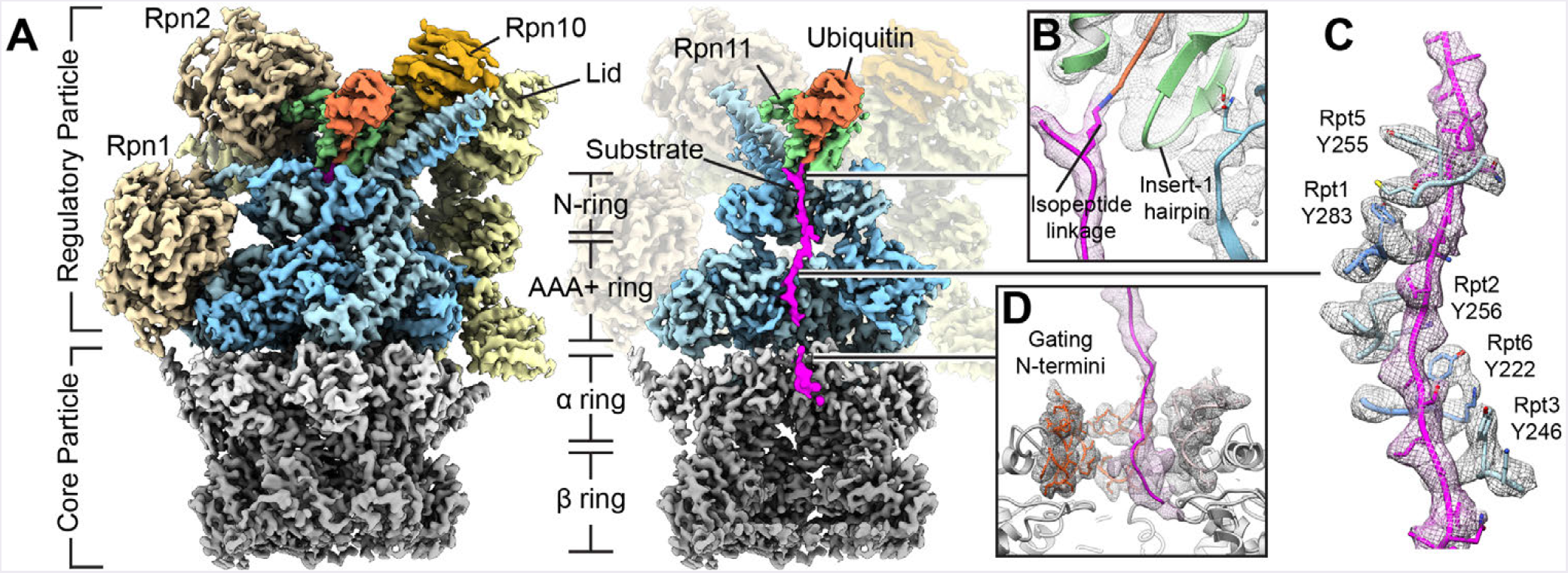
High-resolution structure of the substrate-engaged 26S proteasome. (A) Exterior (left) and cutaway (right) views of the substrate-engaged proteasome cryo-EM reconstruction. The substrate (magenta) is shown extending from the ubiquitin moiety (orange), through the central pore formed by the N-ring and the AAA+ motor (blue), into the gate of the 20S core particle (gray). (B) The isopeptide bond between the substrate lysine and the C-terminus of the ubiquitin moiety is bound in the catalytic groove of Rpn11 (green), with the Insert-1 region in its active, beta-hairpin state that is stabilized by a contact to the N-terminal helix of Rpt5 (blue). (C) The substrate polypeptide is encircled by a spiral-staircase of pore-1-loop tyrosines projecting from the Rpt subunits. (D) Substrate enters the open gate of the core particle. The gating N-termini of α subunits 2, 3, and 4 (red) extend toward the AAA+ motor, forming a hydrophobic collar in conjunction with the N-termini of the other four α subunits (pink).

## Proteasome interactions with the translocating substrate

The stalled proteasome states not only revealed the detailed path of the substrate polypeptide from the RP to the CP, but also resolved the structure of ubiquitin-bound Rpn11 in the context of the 26S holoenzyme(Fig. 1A, Movie 1). The most proximal, substrate-attached ubiquitin moiety of the poly-ubiquitin chain is positioned in the catalytic groove of Rpn11, whose Insert-1 region adopts the same active beta-hairpin conformation previously observed in the crystal structure of the isolated ubiquitin-bound Rpn11/Rpn8 dimer (Fig. 1B, SFig. 3B). While the catalytic Zn^2^+ ion is not visible in the Rpn11 active site, likely due to the treatment with ortho-phenanthroline, the conformations of ubiquitin and Rpn11 match the active, Zn^2^+-containing structure (SFig. 3B), with the addition of an intact isopeptide bond to the substrate lysine.

Upstream (N-terminal) of the ubiquitin-modified lysine, only two amino acids of the substrate were resolved. The orientation of these residues delineates a path near the N-terminal helix of Rpt2 by which substrates may approach the central pore of the proteasome (Fig. 1A), yet to what extent this path outside the N-ring is fixed or substrate-dependent remains unclear. Downstream (C-terminal) of the ubiquitinated lysine, the substrate is confined to the narrow central channel of the Rpt hexamer (Fig. 1A, SFig. 3C-D). An axial view of the RP reveals that the Rpn11 catalytic groove is aligned with the trajectory of substrate translocation through this channel, which follows a straight line from the isopeptide bond into the AAA+ motor (SFig. 3D). This alignment explains how vectorial tugging by the motor can pull ubiquitin directly into the cup-shaped Rpn11 binding site and thus accelerate co-translocational deubiquitination^38^. The active beta-hairpin conformation of Rpn11’ s Insert-1 region thereby seems to be stabilized through additional contacts with Rpt5 at the base of the Rpt4/Rpt5 coiled coil (Fig. 1B). Our structures thus suggest that the translocation stall originates from ubiquitin becoming trapped as it is pulled into the catalytic groove of inactive Rpn11, rather than sterically clashing with the narrow entrance of the N-ring.

Due to the defined stall at the single ubiquitin chain, we were able to reliably model the C-terminally inserted substrate and assign a specific sequence to the polypeptide density within the AAA+ motor (Fig. 1C). The pore-1 loop Tyr and neighboring Lys residues of individual Rpt subunits form a spiral staircase that tightly encircles the substrate, consistent with a translocation mechanism that involves steric interactions with amino-acid side chains of the polypeptide (Fig. 1C). As in many other AAA+ motors, the pore-2 loops are arranged in a second staircase that lies in close proximity to substrate below the pore-1-loop spiral (SFig. 3E). In contrast to the pore-1 loops, the pore-2 loops do not contain bulky residues and may contribute to translocation through interactions of their backbones with the substrate (SFig. 3E), as suggested by defects previously observed for pore-2 loop mutations^20^.

After traversing the AAA+ motor, the substrate enters the gate of the CP (Fig. 1D). Our four cryo-EM structures reveal two gating conformations with distinct RP-CP interactions and arrangements for the N-termini of CP α-subunit (SFig. 4). In all four substrate-bound proteasome structures, the C-termini of HbYX-motif (Hydrophobic-Tyr-anything) containing Rpt subunits (Rpt2, Rpt3, and Rpt5) invariably occupy the intersubunit pockets of the CP α ring, while the pockets for the tails of Rpt1 and Rpt6 vary in occupancy (SFig. 4A). Two of our four proteasome structures show all Rpt tails except for Rpt4 docked into the inter-subunit pockets and consequently a completely open gate, similar to previously described states in substrate-free proteasomes^14,35^ (SFig. 4B-D). In this open conformation, the gating N-termini of α subunits 2, 3, and 4 become directed towards the base subcomplex and interact with the N-termini of the other four α subunits through a conserved Tyr residue (SFig. 4E). This results in the formation of a hydrophobic collar directly beneath the exit from the AAA+ motor (SFig. 4E). The other two structures, which exhibit lower levels of Rpt1- and Rpt6-tail occupancies in the respective α-ring pockets, reveal a partially open gate (SFig. 4C-D). This observation supports a recently proposed model in which cooperative gate opening is driven by the tailFigure 2. Nucleotide-pocket analysis of three sequential substrate-engaged AAA+ motor conformations. (A) Top view of AAA+ motor density maps for three sequential states, named 5D, 5T and 4D, with the substrate-disengaged Rpt subunits indicated by a dashed line. Substrate density (magenta) is shown in the central pore formed by Rpt1 (green), Rpt2 (yellow), Rpt6 (blue), Rpt3 (orange), Rpt4 (red), and Rpt5 (light blue). (B) Close-up views of the Rpt3 nucleotide-binding pocket showing the neighboring Rpt4 providing the Arg finger (top row), and the Rpt5 binding pocket with Arg finger from the neighboring Rpt1 (bottom row). Individual states, left to right, are arranged in the order of motor progression. (C) Measurements of nucleotide-pocket openness colored by nucleotide-bound Rpt subunit. Shown are the distances between the α-carbon of Walker-A Thr and the α-carbon of the neighboring subunit’ s Arg finger (left) or the centroid of α-helix 10 flanking the ISS motif (right). (D) Contact area between the large AAA+ domains of neighboring Rpt subunits.insertion of Rpt1 and Rpt6, after the three HbYX-containing tails are docked^14,35^.

## Distinct nucleotide-states give rise to four ATPase conformations

Our four substrate-engaged proteasome structures show unique motor conformations with one or two disengaged subunits and nucleotide density present in all six ATP-binding pockets (Fig. 2A, S5A-E). To reliably assign nucleotide identities, and thereby establish the progression of the ATP-hydrolysis cycle within the actively hydrolyzing Rpt hexamer, we assessed not only the occupying nucleotide densities, but also the geometries of the ATPase sites, the structural stability of allosteric motifs, and the inter-subunit contact areas (Fig. 2 and Supplementary Table 2). ATP-bound, hydrolysis-competent subunits form a closed pocket with an increased inter-subunit contact area that is characterized by a direct interaction between the γ-phosphate of ATP and the well-resolved Arg fingers of the clockwise neighboring subunit (Figs. 2B-D, S5B-F). In contrast, ADP-bound subunits are more open with a decreased inter-subunit contact area and Arg fingers that are more flexible, as indicated by lower resolvability (Figs. 2B-D, S5B-F). Subunits that are ATP-bound but not yet hydrolysis-competent and subunits directly following ATP hydrolysis show similar, intermediate Arg-finger distances (Figs. 2B-C). To distinguish between these pre- and posthydrolysis states, we assessed the pocket openness by measuring the inter-subunit contact area or the distance between the conserved Walker-A-motif Thr and the helix preceding the inter-subunit signaling (ISS) motif of the neighboring subunit (Figs. 2C-D, S5E)^26,33^. Our analyses revealed a continuum of nucleotide states within the Rpt hexamers, ranging from ATP-bound partially open pockets with semi-engaged Arg fingers (hydrolysis-incompetent), ATP-bound closed pockets with fully engaged Arg fingers (hydrolysis-competent), ADP-bound closed pockets with disengaged Arg fingers (posthydrolysis), and ADP-bound open pockets with disengaged Arg fingers (pre-nucleotide exchange) (Fig. 2B). Individual Rpt subunits show a progression through discrete nucleotide states around the hexameric ring (Fig. 3A), indicating that each reconstruction reflects a unique snapshot of the proteasomal AAA+ motor during the ATPase cycle, and that Rpts likely progress sequentially through this cycle. Our structures thus provide the unique ability to correlate the distinct vertical registers and nucleotide states of Rpt subunits, and analyze the coupling of individual ATPase steps with each other and with the mechanical translocation of substrate.

**Figure 2.**
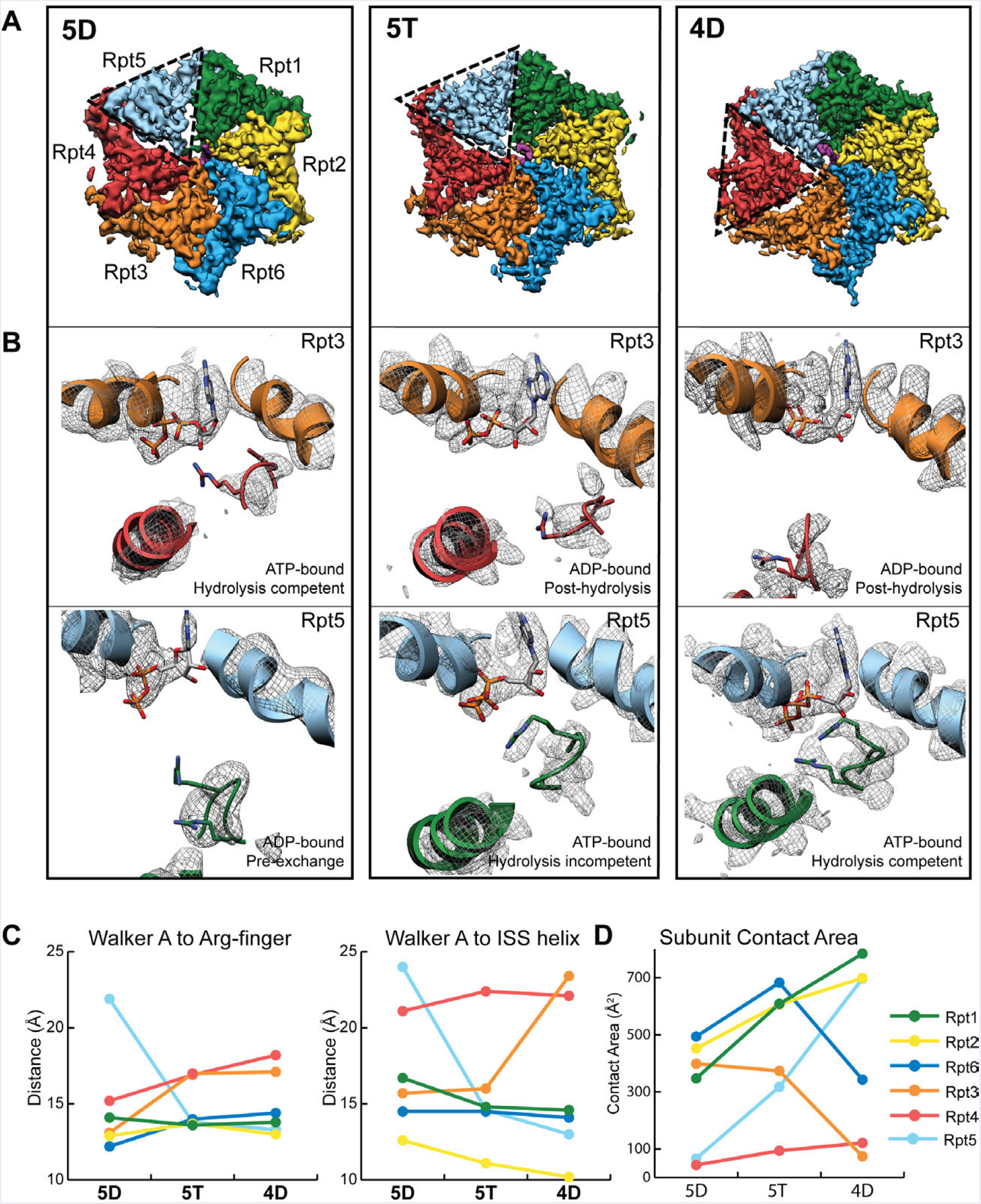
Nucleotide-pocket analysis of three sequential substrate-engaged AAA+ motor conformations. (A) Top view of AAA+ motor density maps for three sequential states, named 5D, 5T and 4D, with the substrate-disengaged Rpt subunits indicated by a dashed line. Substrate density (magenta) is shown in the central pore formed by Rpt1 (green), Rpt2 (yellow), Rpt6 (blue), Rpt3 (orange), Rpt4 (red), and Rpt5 (light blue). (B) Close-up views of the Rpt3 nucleotide-binding pocket showing the neighboring Rpt4 providing the Arg finger (top row), and the Rpt5 binding pocket with Arg finger from the neighboring Rpt1 (bottom row). Individual states, left to right, are arranged in the order of motor progression. (C) Measurements of nucleotide-pocket openness colored by nucleotide-bound Rpt subunit. Shown are the distances between the a-carbon of Walker-A Thr and the a-carbon of the neighboring subunit’s Arg finger (left) or the centroid of a-helix 10 flanking the ISS motif (right). (D) Contact area between the large AAA+ domains of neighboring Rpt subunits.

**Figure 3.**
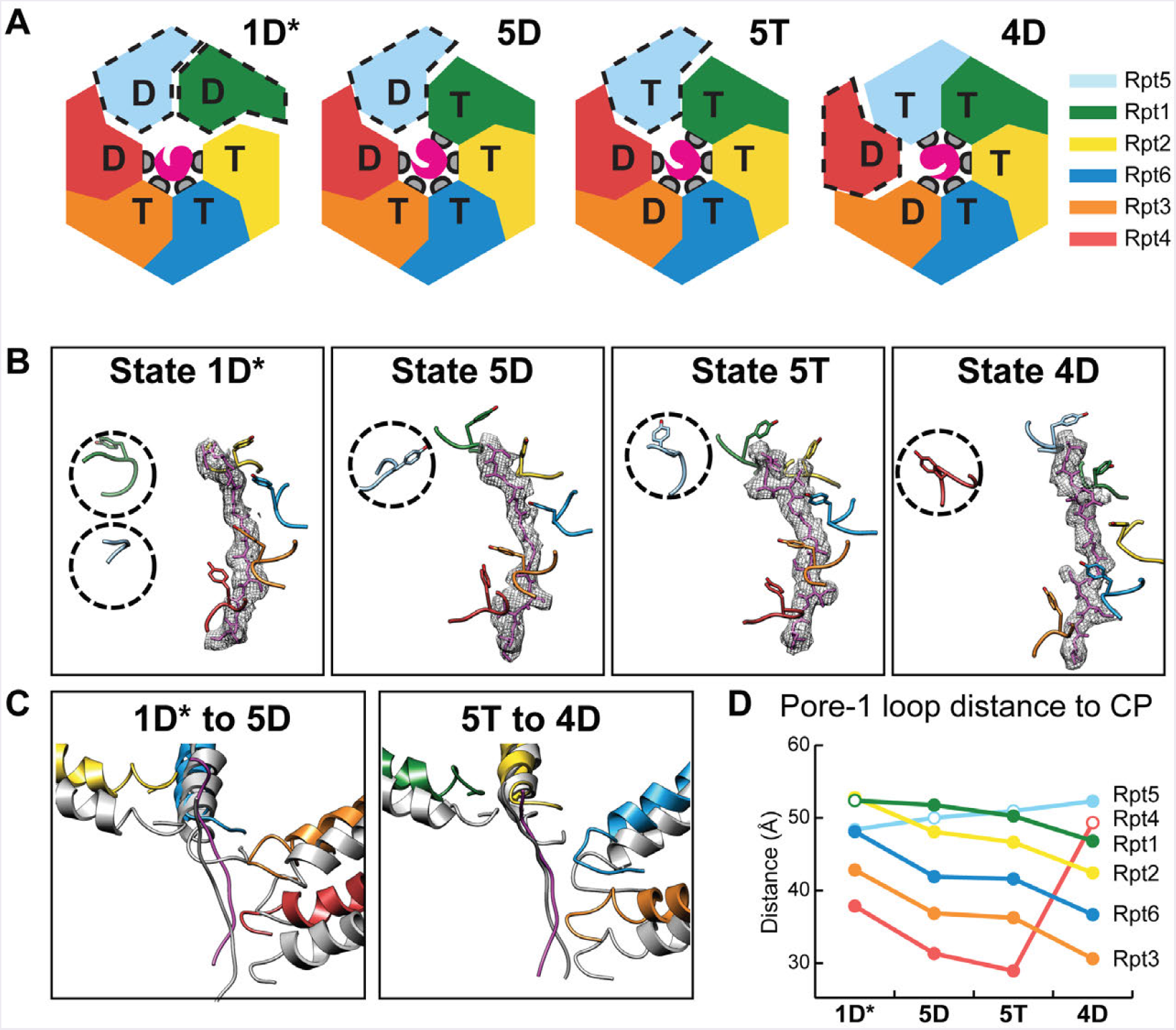
Pore-1 loop tyrosines define three distinct spiral-staircase conformations of the AAA+ motor. (A) Summary of nucleotide states and staircase arrangement in the 1D*, 5D, 5T and 4D states. Coloring of the motor subunits Rpt1 (green), Rpt2 (yellow), Rpt6 (blue), Rpt3 (orange), Rpt4 (red), and Rpt5 (light blue) is consistent throughout the figure. Pore-1 loop contacts (gray) with substrate (magenta) are not present in the disengaged subunits (dashed outline). (B) Pore-1 loop Tyr staircases for each of the substrate-bound states. Substrate polypeptide (mesh) is encircled by four or five engaged pore-1 loops in each state. Disengaged pore-1loops are indicated by dashed circles. (C) Vertical movement of substrate-engaged pore-1 loops is observed during motor transition from 1D* to 5D and 5T to 4D states. The lower state (5D and 4D, respectively) is shown in gray. (D) Plot of distances between the a-carbon of the pore-1 loop tyrosines to the plane of the core particle’s gate. Filled circles represent substrate-engaged pore loops, open circles indicate disengaged pore loops

## Sequential motor states reveal the mechanism for ATP-hydrolysis-driven substrate translocation

The current hand-over-hand translocation model for hexameric AAA+ ATPases specifies that the subunits encircle and interact with substrate in a staircase-like organization, with the exception of the sixth subunit (often referred to as the “ seam subunit”) that is displaced from the substrate and positioned between the lowest and highest subunits of the staircase^25-30^. In accordance with this established configuration for substrate translocation, we see that in all substrate-engaged proteasome states, the Rpt subunits interact with the substrate through their pore-1 loops in a spiral-staircase arrangement, with the pore-2 loops forming a similar staircase underneath (Figs. S6A-C). A characteristic seam is observed along the interface between the highest substrate-engaged subunit of the staircase and the neighboring substrate-disengaged subunit (Figs. 2A, 3A-B, S5A). We name our four proteasome states based on the identity and the nucleotide state of this substrate-disengaged “ seam” subunit: Rpt1-ADP; Rpt5-ADP; Rpt5-ATP; and Rpt4-ADP, as 1D* 5D; 5T; and4D, respectively (Fig. 3A).

In these four conformations, three different Rpt subunits occupy the upper-most substrate-bound position of the staircase. In 1D*, Rpt2 is in the top substrate-bound position with the seam subunit Rpt1 displaced from the substrate (Fig. 3B). Unexpectedly, Rpt5 is also disengaged in this conformation, indicating that 1D* may represent an off-pathway ATPase configuration (see discussion below).

We posit that the remaining three conformations—5D, 5T, and 4D—represent consecutive states, whose transitions include a nucleotide exchange, a hydrolysis event, and a translocation step (Fig. 3, Movie 2). In 5D and 5T, the vertical staircase register is shifted by one subunit in the counterclockwise direction compared to 1D*, such that Rpt1 assumes the uppermost substrate-bound position, while the other subunits move downwards, and only Rpt5 is substrate-disengaged (Fig. 3B). During the 5D-to-5T transition, the staircase arrangement of Rpt subunits remains largely the same (Fig. 3D, SFig. 6D), but the density for the Rpt5-bound nucleotide changes concomitantly with a substantial closure of the binding pocket that brings the Arg fingers of Rpt1 into close proximity (Fig. 2B), which is consistent with an exchange of ADP for ATP. This exchange and the resulting shift of Rpt5 towards the central pore likely primes this subunit by allosterically positioning the pore loops for substrate engagement in the subsequent 4D state (Fig. 3B). Nucleotide exchange in the disengaged Rpt thus appears prerequisite for substrate binding at the top of the spiral staircase, which agrees with the highest substrate-contacting subunit always being ATP-bound (Fig. 3A). Importantly, concurrent with ATP binding to Rpt5, Rpt3 hydrolyzes ATP during the 5D-to-5T transition, as indicated by correlative changes in the Rpt3-bound nucleotide density and the disengagement of the neighboring Arg finger (Figs. 2B, 2C). Neither the nucleotide exchange nor the hydrolysis event cause significant conformational changes in the Rpt hexamer but represent the trigger for the most striking rearrangement of the mechanochemical cycle in the subsequent transition to 4D.

During this 5T-to-4D transition, we observe an opening of the Rpt3 nucleotide-binding pocket and a disruption of the inter-subunit interactions with the neighboring Rpt4 (Figs. 2B, 2D). Rpt4 separates from Rpt3, disengages from substrate, and moves from the bottom of the staircase out and upwards, which is likely driven by the topologically closed ring architecture of the Rpt hexamer. At the same time, the ATP-bound Rpt5 at the top of the staircase moves to a more central positon and binds substrate (Fig. 3B), while the substrate-engaged subunits Rpt1, Rpt2, Rpt6, and Rpt3 move as a rigid body downwards by one register and translocate the substrate toward the CP gate (Figs. 3C-D, Movie 2).

Even though we do not detect concrete nucleotide-density changes during this transition, we can postulate based on the preceding ATP-hydrolysis event in Rpt3 and the subsequent opening of its pocket that phosphate release from Rpt3 is responsible for the disruption of inter-subunit interactions with Rpt4 and the consequent conformational changes of the entire ATPase ring. This model is consistent with our observations that the penultimate subunit in the staircase exhibits a completely or partially closed pocket in all proteasome conformations, whereas the lowest substrate-engaged subunit is always ADP-bound with an open pocket (Figs. 3A, 2C-D, S5B-E). Furthermore, it agrees with previous single-molecule data on the homohexameric ClpX ATPase, suggesting that phosphate release represents the force-generating step of the ATPase cycle^39^. Similar to the coordinated nucleotide-exchange and ATP hydrolysis steps in the previous 5D-to-5T transition, the disruption of the Rpt3-Rpt4 interface through potential phosphate release, the substrate-engagement by Rpt5, and the movement of a four-subunit rigid body for substrate translocation appear to be interdependent and tightly coupled during the 5T-to-4D transition.

For the ATP-hydrolysis and substrate-translocation cycles, our findings suggest that a particular subunit binds ATP and engages substrate at the uppermost position, hydrolyzes ATP when at the penultimate position of the staircase, releases phosphate as it moves to the bottom of the ring, before disengaging from substrate in the next step (Fig. 4A). AAA+-motor movements and substrate translocation would thus be powered by sequential ATP hydrolysis and phosphate release as each Rpt transitions to the bottom of the staircase. This model, wherein all Rpt subunits cycle through the various conformational states and vertical registers of the staircase, is further supported by previously determined structures of the 26S proteasome in the absence of substrate (see below).

**Figure 4.**
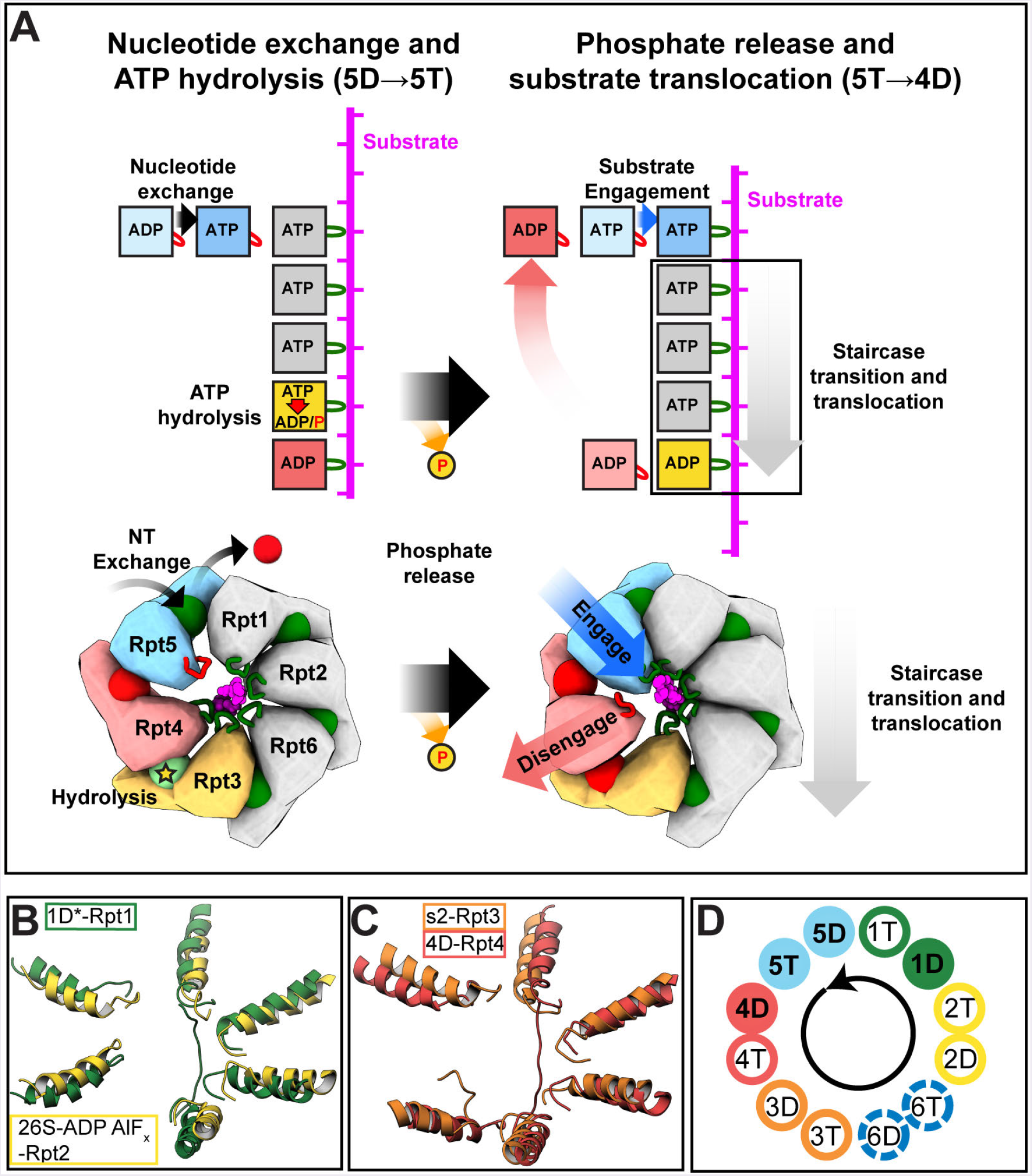
Coordinated ATP-hydrolysis and substrate-translocation cycles of the proteasome.(A) Model for the coordination of ATP-hydrolysis steps and their coupling to substrate translocation. Nucleotide exchange and ATP hydrolysis occur simultaneously in the substrate-disengaged (blue) and penultimate subunit (orange) of the staircase, respectively, with no major conformational changes of the motor. Subsequent phosphate release from the penultimate subunit leads to the displacement of the bottom subunit (red), substrate-engagement by the top subunit (blue), and downward movement of a four-subunit rigid body (boxed) to translocate substrate. (B) Spiral staircases of 1D* (dark green) and the 26S-ADP-AlF_x_ (yellow; PDB:5wvk^35^), rotated to overlay the disengaged subunits (1D*-Rpt1 and 26S-ADP-AlF_x_-Rpt2) and aligned by the pore-1 loop helices. (C) Spiral staircases of the s2 state (orange; PDB:5mpa^33^) and the 4D state, rotated to overlay the disengaged subunits (s2-Rpt3 and 4D-Rpt4) and aligned by the pore-1 loop helices. (D) Schematic of the possible progression of proteasome states, colored by the disengaged subunit, with our observed staircase states indicated by filled circles, staircases from substrate-free proteasome states indicated by open circles, and potential additional states represented by dashed circles.

Our observation of four substrate-engaged subunits moving as a rigid body to translocate substrate in response to ATP hydrolysis and phosphate release is consistent with previous biochemical studies of the ClpX ATPase, which indicated that several subunits interact synergistically with substrate, allowing even a pore-1-loop-deficient subunit to drive translocation^40,41^. The rigid-body movement of four Rpts vertically advances the engaged pore-1-loop Tyr by about 6 Å (Fig. 3C-D), suggesting a fundamental step size of two amino acids per hydrolyzed ATP for proteasomal substrate translocation.

We do not observe a vertical movement of substrate due to the defined stall of translocation upon Rpn11 inhibition, and all our proteasome conformations show largely the same stretch of polypeptide in the central channel. Nevertheless, the substrate responds to staircase rearrangements with lateral movements in the ATPase channel, shifting towards the engaged pore-1 loops and away from the disengaged subunits (SFig. 6A). The substrate backbone follows the spiral-staircase arrangement of pore loops, rather than traversing the motor in a straight vertical path, and its lateral position in the channel rotates counterclockwise around the hexamer as the Rpts progress through the various nucleotide states (SFig. 6A).

## Additional states of the proteasomal ATPase cycle

While 5D, 5T, and 4D each contain four ATP-bound and two ADP-bound subunits, 1D* shows three ATP-bound and three ADP-bound subunits (Fig. 3A), and we interpret this conformation as an alternate, potentially off-pathway version of a 1D state. A comparison of the Rpt subunit organization in 1D* with those in the 5D, 5T, and 4D states as well as other proteasome- and AAA+ motor structures (Figs. S7A-B), suggests that Rpt5 has prematurely released from substrate at the bottom of the spiral after opening of the Rpt4 nucleotide-binding pocket, and hence both Rpt5 and Rpt1 are disengaged (Figs. 3B, S7A). Notably, the Rpt5 pore-1 loop is divergent from the other Rpts, containing a conserved Met rather than a Lys^42^, which could result in weaker substrate interactions, especially when translocation is stalled, and thus contribute to the premature disengagement in the 1D* state. Conversely, the 1D* state can be explained by failed nucleotide exchange in Rpt1 at the top of the spiral, which prevented substrate engagement and the consequent rearrangement of the staircase to the 5D state. Interestingly, the previously described s3 conformation of the substrate-free proteasome shows the expected staircase arrangement of 1D, with Rpt5 remaining in the lowest position of the staircase (SFig. 7B)^33^.

Our three distinct spiral-staircase states offer a view of the discrete events leading to a complete step of hydrolysis-driven substrate translocation, yet the current hand-over-hand model requires that every Rpt subunit cycles through all the states. Previous biochemical studies have indicated that ATP hydrolysis in almost all Rpt subunits contribute to substrate engagement and translocation^20,43,44^. Therefore, the proteasome conformations described here likely represent only a subset of states that may be complemented by the corresponding 4T, 3D, 3T, 6D, 6T, 2D, and 2T states, as well as an additional 1T state between 1D and 5D (Figs. 4B-D, S7), to complete the ATP-hydrolysis and substrate-translocation cycles of the proteasome.

Indeed, several additional staircases have been previously observed for the proteasome, albeit in the absence of substrate and induced by non-hydrolyzable ATP analogs, which hampered robust derivations about mechanochemical coupling or the ATPase cycle. Their overall similarity to our Rpt staircase arrangements is sufficient to designate specific spiral staircases. Some of these states (e.g. the ADP-AlF_x_-bound and s2 states) were regarded as unlikely processing conformations of the AAA+ motor, as they were associated with a partially open CP gate^14,33,34^. However, two of our engaged states similarly contain only partially open gates, yet clearly show substrate being threaded through the central channel to the CP gate (SFig. 4). This indicates that a fully open gate is not required for every step of substrate translocation, but its openness may vary depending on the state of the Rpt staircase and corresponding allosteric subtleties in Rpt-tail interactions with the CP. More accurate criteria for a processing motor state are the coaxial alignment of the AAA+ motor with CP, the rotation of the lid subcomplex, and the presence of rigid bodies formed between the large AAA subdomain of one subunit and the small AAA subdomain of its neighbor in all but the substrate-disengaged Rpts. Based on these criteria and their staircase orientation, the s2 and recently described s5 states^14^ would represent 3D or 3T states, and the ADP-AlFx-bound proteasome conformation^34^ resembles a putative 2D* or 2T* state, with two substrate-disengaged subunits similar to 1D* (Figs. 4B-C). The s4 and SD2 states, which had previously been proposed as potential processing states^33,35^, show overall staircase similarities with our substrate-engaged 4D and 5D states, respectively, even though some of their Rpts are distorted as a likely consequence of inhibited ATP hydrolysis and the absence of substrate (Figs. S7C-D).

Our substrate-engaged proteasome states hence provide a structural context for previously described ATP-analog-bound conformations, enabling us to approximate all possible Rpt staircases, except for the hypothetical 6D and 6T states, and thus support our model of a sequential hand-over-hand mechanism wherein each Rpt transitions through the ATP-hydrolysis and substrate-translocation cycles (Fig. 4D). Why the protea-some in the presence of substrate preferentially adopts only the 1D*, 5D, 5T, and 4D states remains unclear, but it is possible that these conformations are favored because of the substrate stall or represent dwell states that are longer lived due to intrinsic differences in the rates for nucleotide binding, hydrolysis, or exchange. Importantly, however, the consecutive 5D, 5T, and 4D states are sufficient to provide us with a complete picture of subunit transitions during the ATPase cycle and substrate translocation.

## Conclusion

In this study, we elucidated the first structures of the substrate-engaged 26S proteasome that answer many of the outstanding questions regarding proteasomal degradation and the general mechanism by which AAA+ translocases process their substrates. Resolving the ATP-hydrolyzing AAA+ motor in distinct conformations with a stalled, ubiquitinated substrate polypeptide provides unprecedented insights into the coordination of ATP-hydrolysis steps, their mechanochemical coupling with translocation, and the mechanism underlying co-translocational deubiquitination.

Our structures show that Rpn11 inhibition causes a substrate stall in which the proximal moiety of the substrate-attached poly-ubiquitin chain remains functionally bound within the Rpn11 catalytic groove. This trapped state revealed a linear alignment of the scissile isopeptide bond at the Rpn11 active site with the translocation trajectory of the substrate polypeptide through the AAA+ motor (Fig. 1). We conclude that during normal degradation, ubiquitin modifications are pulled directly into the Rpn11 catalytic groove, and that further substrate translocation is arrested until the isopeptide has been cleaved. This ubiquitin-capture mechanism explains how Rpn11 functions as a gate keeper to efficiently remove all ubiquitin modifications on a substrate, and how deubiquitination can be accelerated by mechanical pulling of the AAA+ motor on the substrate polypeptide^38^.

The Rpt subunit conformations we detected in the substrate-engaged, actively ATP-hydrolyzing proteasomes validates the mechanistic relevance of the staircase arrangements seen in structures for a variety of related, substrate-bound homohexameric AAA+ motors, whose stabilization required nucleotide analogs or ATPase-inactivating mutations^25-29^. Mechanistic models based on these previous homohexamer structures rely on the assumption that individual snapshots are representative of all possible spiral-staircase conformations. Furthermore, the coordination of ATP-hydrolysis steps and their coupling to the conformational changes that drive substrate translocation through the hexamer remained elusive due to the lack of intermediate or directly consecutive states. Here we resolved three sequential states of the Rpt heterohexamer, which provides a model for the inter-subunit coordination during nucleotide exchange, ATP hydrolysis, and phosphate release within the AAA+ motor and how these events are mechanochemically coupled to substrate translocation.

Consistent with other substrate-bound AAA+ ATPase structures, our proteasome motor adopts staircase arrangements encircling an unfolded polypeptide substrate, with one substrate-disengaged subunit^25-29,45^ (Fig. 3). We observed that four of the substrate-engaged subunits are ATP-bound, whereas the subunit at the bottom of the staircase and the disengaged subunit are ADP-bound (Fig. 3A). Our structures suggest that nucleotide exchange primes the disengaged subunit for substrate binding at the top of the staircase, and that this exchange occurs concomitantly with ATP hydrolysis in the fourth substrate-engaged subunit of the staircase. Both steps of the ATPase cycle are associated with only subtle allosteric rearrangements, whereas the entire ATPase hexamer undergoes major conformational changes during the subsequent transition that appears to be linked to phosphate release from the post-hydrolysis, fourth subunit of the staircase. These rearrangements include the displacement of the bottom ADP-bound subunit, substrate-binding of the previously disengaged subunit at the top of the staircase, and the downward movement of the remaining four substrate-engaged subunits as a rigid body. It seems that all these processes must happen in a coordinated fashion for substrate translocation to occur.

This translocation mechanism is thus reminiscent of a six-subunit conveyor belt, in which a four-subunit rigid body grips the substrate and moves downwards as the bottom-most subunit disengages and the top-most subunit re-engages substrate (Fig. 4A). The coordinated gripping by pore-loops of four subunits, which are stabilized by ATP-bound, closed interfaces, likely enables higher pulling forces and reduced slippage, consistent with previous biochemical studies of the ClpX motor^40,41^. Similar conveyor-belt mechanisms have been proposed previously for AAA+ protein translocases as well as DNA and RNA helicases^22,23,26-29,45^, yet our structures notably clarify the precise movement of ATPase subunits and their coordination with individual steps of the ATPase cycle.

Given the strong structural and functional similarities between the proteasomal Rpt hexamer and other AAA+ motors, we hypothesize that our proposed model for ATP-hydrolysis-coupled substrate translocation applies to hexameric AAA+ translocases in general. Furthermore, the consecutive ATPase states that we observe, together with equivalent staircases in previous substrate-free proteasome structures, suggest a sequential progression of individual Rpt subunits through the ATPase cycle, rather than a burst mechanism, where several subunits hydrolyze in rapid succession prior to nucleotide exchange, as proposed for the ClpX motor based on singlemolecule measurements^46,47^. The dwell times for the hetero-hexamer progressing through the various stages of the ATPase cycle may nonetheless differ for different Rpts, which could explain why we observe only four of the potential staircase states. Additional biophysical analyses, for instance of nucleotide on and off rates for individual Rpts, will therefore be required to further elucidate the temporal dynamics of the proteasomal ATPase motor and the timing of subunit transitions for mechanical substrate translocation.

## Materials and Methods

### Sample Preparation

Purification of proteasome Holoenzyme: 26S proteasomes were purified from strain YYS40 (MATa leu2-3,112 trp1-1 can1-100 ura3-1ade2-1 his3-11,15 RPN11::RPN11-3XFLAG (HIS3)^48^ as previously described^49^. Briefly, frozen yeast paste from saturated cultures was lysed in a Spex SamplePrep 6875 Freezer/Mill, and cell powder was resuspended in 60 mM HEPES, pH 7.6, 20 mM NaCl, 20 mM KCl, 8 mM MgCl2, 2.5% glycerol, 0.2% NP-40, and ATP regeneration mix (5 mM ATP, 0.03 mg/mL creatine kinase, 16 mM creatine phosphate). Proteasomes were batch-bound to anti-FLAG M2 Affinity Gel (Millipore Sigma), washed with Wash Buffer (60 mM HEPES, pH 7.6, 20 mM NaCl, 20 mM KCl, 8 mM MgCl2, 2.5% glycerol, 5mM ATP), eluted with 3XFLAG peptide, and further separated by Size Exclusion Chromatography using a Superose 6 Increase column in 60 mM HEPES, pH 7.6, 20 mM NaCl, 20 mM KCl, 10 mM MgCl2, 2.5% glycerol, and 1 mM ATP.

Preparation of ubiquitinated model substrate: A model substrate consisting of an N-terminal Cys, lysine-less titin-I27V15P, a single-lysine-containing sequence derived from an N-terminal fragment of *Sphaerechinus granularis* cyclinB (residues 22-42, with Lys to Ala substitutions), a Rsp5 recognition motif (PPPY), and 6X His-tag, was purified after expression in E. coli BL21-Star by standard methods. Briefly, In Terrific Broth, protein expression was induced with IPTG at OD600 =1.2-1.5 for 5 hours at 30°C. Cells were harvested by centrifugation, resuspended in chilled Lysis Buffer (60 mM HEPES, pH 7.6 100 mM NaCl, 100 mM KCl, 15 mM Imidazole) and lysed by sonication. Following clarification by centrifugation at 20,000 x g, the protein was purified using Ni-NTA affinity chromatography. The substrate was fluorescently labeled using 5-Fluoroscein maleimide at pH 7.2 for 3 hours at room temperature and quenched with DTT. Free dye was separated from the substrate by Size Exclusion Chromatography with a Superdex 200 column (GE Healthcare), buffer exchanging the substrate into 60 mM HEPES, pH 7.6, 20 mM NaCl, 20 mM KCl, 10 mM MgCl2, 2.5% glycerol.

The substrate at final substrate concentration of 50 μM was modified with long, K63-linked ubiquitin chains using 5 μM Mus musculus Uba1, 5 μM *S. cerevisiae* Ubc1, 20 μM S. cerevisiae Rsp5ΔWW^20,50^, and 2 mM S. cerevisiae ubiquitin, in 60 mM HEPES, pH 7.6, 20 mM NaCl, 20 mM KCl, 10 mM MgCl2, 2.5% glycerol, and 15 mM ATP for 3 hours at 25°C, followed by incubation overnight at 4°C.

### ATPase Assay

Proteasome ATPase activity was monitored using a spectrophotometric assay that couples regeneration of hydrolyzed ATP to the oxidation of NADH^51^. Reactions contained a final concentration of 150 nM 26S proteasome that had been preincubated with ortho-phenanthroline and ATPase mix for 5 minutes on ice or mock treated before bringing the sample to 25°C and adding FAM-labeled ubiquitinated substrate to a final concentration of 3 μ M and ortho-phenanthroline to a final concentration of 3 mM. Absorbance at 340 nm was measured for 10 minutes with 12-second intervals in a 384-well plate (Corning) using a Biotek Synergy Neo2 plate reader. Reactions were done in 60 mM HEPES, pH 7.6, 20 mM NaCl, 20 mM KCl, 10 mM MgCl_2_, 2.5% glycerol 1mM TCEP, and 1 X ATPase mix (5 mM ATP, 3 U ml^-1^ pyruvate kinase, 3 U ml^-1^ lactate dehydrogenase, 1 mM NADH, and 7.5 mM phosphoenol pyruvate).

### Gel-based monitoring of proteasome degradation activity

Proteasome was pre-incubated as described in the ATPase assay. 10 minutes after the addition of ubiquitinated substrate, samples were quenched by the addition of 2% SDS and separated on a 4-20% gradient Tris-Glycine SDS-PAGE gel (BioRad). Fluorescence at 530nm from the FAM-labeled substrate was measured using a BioRad ChemiDoc MP imager. Total protein was imaged by staining gels with Colloidal Coomassie Brilliant Blue following fluorescence imaging.

### Grid preparation for cryo-electron microscopy

26S proteasomes were diluted to a concentration of 20 μM in a solution containing 20 mM HEPES, pH 7.6, 25 mM NaCl, 25 mM KCl, 10 mM MgCl2, 1 mM TCEP, 5 mM ATP, 0.05% NP-40, an ATP regeneration system, and 6 mM ortho-phenanthroline. This solution was mixed with an equal volume of 50 μM ubiquitinated model substrate. Three microliters of the holoenzyme-substrate solution were immediately applied to R2/2 400-mesh grids (Quantifoil) that had been plasma treated for 20 seconds using a glow discharger (Electron Microscopy Sciences) operated under atmospheric gases. The grids were manually blotted to near dryness with Whatman No. 1 filter paper inside a cold room (4°C) and gravity plunged into liquid ethane using a home-built system.

### Data collection and image processing

Cryo-EM data were acquired using the Leginon software for automated data acquisition^52^ and a Titan Krios (Thermo Fisher) equipped with a K2 Summit (Gatan) direct electron detector in counting mode (Table 1). Movies were collected by navigating to the center of a hole and sequentially image shifting to ten targets situated at the periphery of the 2 μm hole (SFig. 2F). To maximize the number of targets per hole, a nanoprobe beam of 597 nm in diameter was utilized. This resulted in a total acquisition of 11,656 movies at an approximate rate of 2200 movies per day. Movies were recorded at a nominal magnification of 29000 × (1.03 Å magnified pixel size) and composed of 25 frames (250 ms per frame, ~50 e” /Å^2^ per movie). Movie collection was guided by real-time assessment of image and vitrified sample quality using the Appion image-processing software^53^. Frame alignment and dose weighting were performed in real-time using UCSF Motioncor2^54^. CTF estimation on aligned, unweighted, micrographs was performed with Gctf^55^.

All data post-processing steps were conducted in RELION 2.1^56,57^. Holoenzyme particles were picked using s1 proteasome templates generated from 2D class averages obtained from a prior cryo-EM experiment. This resulted in 579,361 particle picks that were extracted (660×660 pixels) and downsampled (110x100 pixels) for reference-free 2D classification. 298,997 particles, belonging to the 2D classes demonstrating features characteristic of secondary structural elements, were subjected to 3D refinement and subsequent 3D classification (k=10). A 3D template of an s1 proteasome was utilized to guide the initial 3D refinement and 3D classification, which ensured that s4-like or substrate-bound reconstructions did not arise from template bias. 238,828 particles corresponding to 3D classes without artefactual features were chosen for further data processing. To minimize the detrimental effects of the holoenzyme’ s pseudo-symmetry (C2) on resolution, the raw holoenzyme particles were C2 symmetry expanded, 3D refined, and a python script was used to determine the x,y coordinates corresponding to the center of the ATPase within the regulatory particles (RP). In this way, the RPs at each end of every core particle were re-extracted to serve as individual asymmetric units without down-sampling. A round of reference-free 2D classification enabled us to remove the ends of core particles that lacked a regulatory particle. This combined expansion and classification approach netted 380,011 unique RP-containing particles.

We performed 3D classification on the RP dataset and isolated 242,980 particles whose parent 3D class exhibited a globular ubiquitin-shaped density in the periphery of the Rpn11 active site. Further classification aimed to identify substrate in the central pore of the proteasome. To accomplish this, a soft mask encompassing the AAA+ motor was used to exclude the rest of the proteasome for 3D classification and 3D refinement. This resulted in four distinct AAA+ motor reconstructions containing density attributed to substrate in the central pore with nominal resolutions ranging from 3.9 to 4.7 Å (SFig. 2). To further increase map quality outside the AAA+ motor, the global maps corresponding to each AAA+ motor were subdivided into 12 regions for focused 3D refinement, and a composite map consisting of all 12 focused 3D refinements was then generated for each reconstruction to facilitate atomic model building (SFig. 2E).

### Atomic model building

All atomic models were built using the s4 proteasome model (PDB ID: 5MPC^33^) as a template. The initial template’ s subunits were individually rigid body fit into each of the four EM reconstructions (1D, 5D, 5T, and 4D) with “ Fit in Map” function of Chimera^58^. The docked templates were then subjected to one cycle of morphing and simulated annealing in PHENIX, followed by a total of 10 real-space refinement macrocycles utilizing atomic displacement parameters, secondary structure restraints (xsdssp), local grid searches, and global minimization^59^. After automated PHENIX refinement, manual real-space refinement was performed in Coot^60^. Residue side chains without attributable density were truncated at the α-carbon, ions were removed, and atoms corresponding to the β-ring, N-ring, and RP lid were removed in the 1D, 5D, and 5T models due to redundancy and to accelerate refinement. For the 4D state, a similar approach was followed, but atoms corresponding to the N-ring were not removed to facilitate template-based (PDB ID: 5U4P^38^) modeling of ubiquitin at the Rpn11 active site. Isopeptide bond-length restraints were manually created and implemented during PHENIX refinement^59^. Multiple rounds of real-space refinement in PHENIX (5 macro cycles, no morphing, no simulated annealing) and Coot were performed to address geometric and steric discrepancies identified by the RCSB PDB validation server and MolProbity^59-61^. All images were generated using UCSF Chimera^58^ and ChimeraX^62^.

## Acknowledgments

We thank B.M. Gardner, J.A.M. Bard, A.L. Yokom, and C. Puchades for comments and critical evaluation of the manuscript, and all members of the Martin and Lander labs for discussions, suggestions, and support. We are grateful to R.J. Beckwith for cloning the model substrate. All cryo-EM data were collected at the Scripps Research (SR) electron microscopy facility. We thank B. Anderson for microscope support and J.C. Ducom at SR High Performance Computing facility for computational support. S.N.G. is a Howard Hughes Medical Institute Fellow of the Damon Runyon Cancer Research Foundation, DRG-2342-18. G.C.L is supported as a Pew Scholar in the Biomedical Sciences by the Pew Charitable Trusts. A.M. is an investigator of the Howard Hughes Medical Institute. This work was funded by the National Institutes of Health (DP2-EB020402 to G.C.L. and R01-GM094497 to A.M.) and the Howard Hughes Medical Institute (A.M.). Computational analyses of EM data were performed using shared instrumentation at SR funded by NIH S10OD021634.

## Data and materials availability

Coordinates for the 1D*, 5D, 5T, and 4D states have been deposited in the Protein Data Bank under accession codes 6EF0, 6EF1, 6EF2, and 6EF3, respectively. The associated cryo-EM reconstructions have been deposited to the Electron Microscopy Data Bank under accession codes: EMD-9042, EMD-9043, EMD-9044, EMD-9045.

---

**Supplementary Figure 1.**
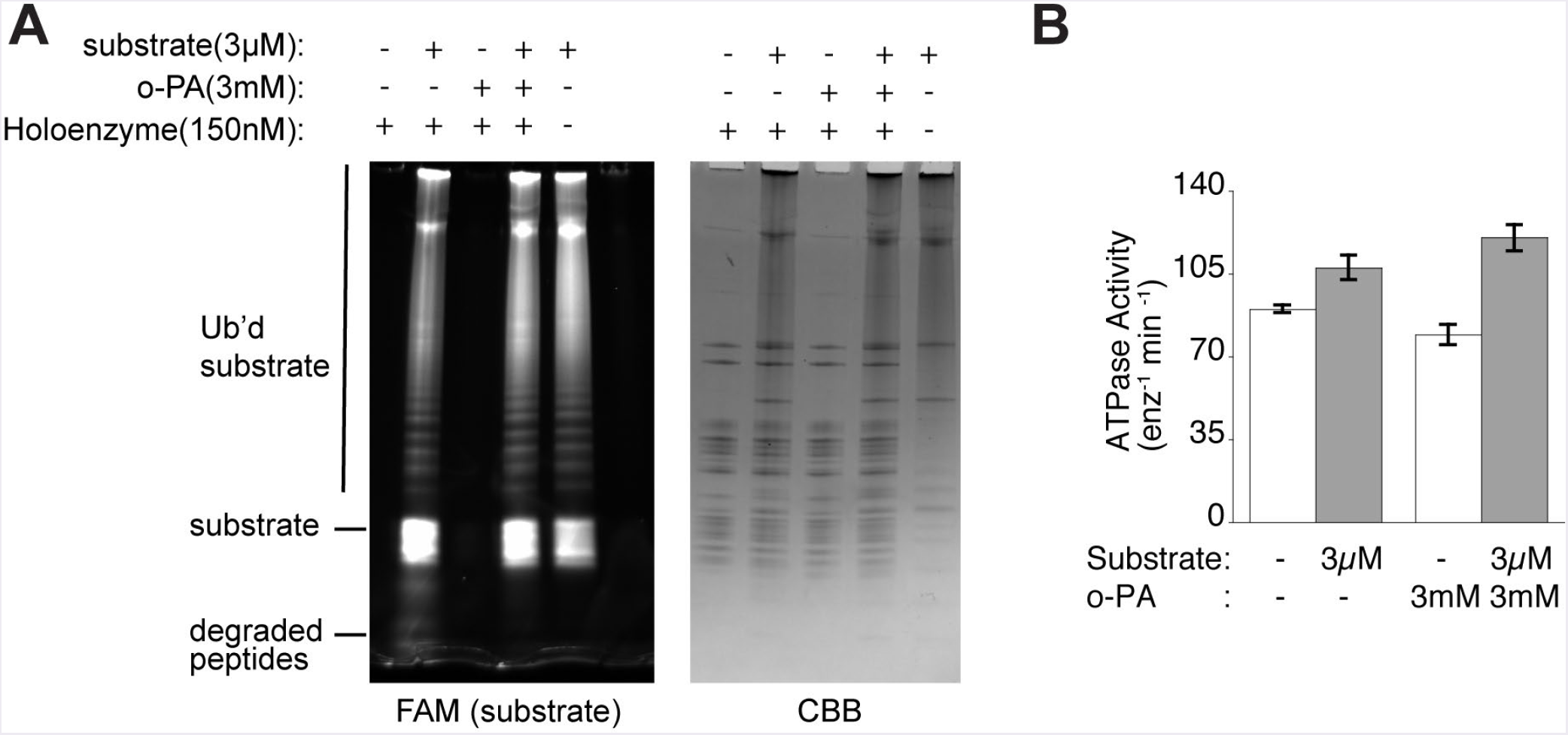
Ortho-phenanthroline-stalled proteasomes do not degrade the substrate but maintain substrate-mediated stimulation of ATPase activity. (A) SDS-PAGE analysis of degradation endpoints after incubation of substrate with ortho-phenanthroline (o-PA) treated or untreated proteasomes. Fluorescence emission (530 nm) is shown for the N-terminally FAM-labeled substrate (left), and total protein is shown by staining with colloidal coomassie brilliant blue (CBB) (right). (B) Proteasomal ATPase activity in the presence or absence of substrate and ortho-phenanthroline (o-PA). Columns represent the mean of three technical replicates with standard error (s.e.m) shown as bars.

**Supplementary Figure 2A.**
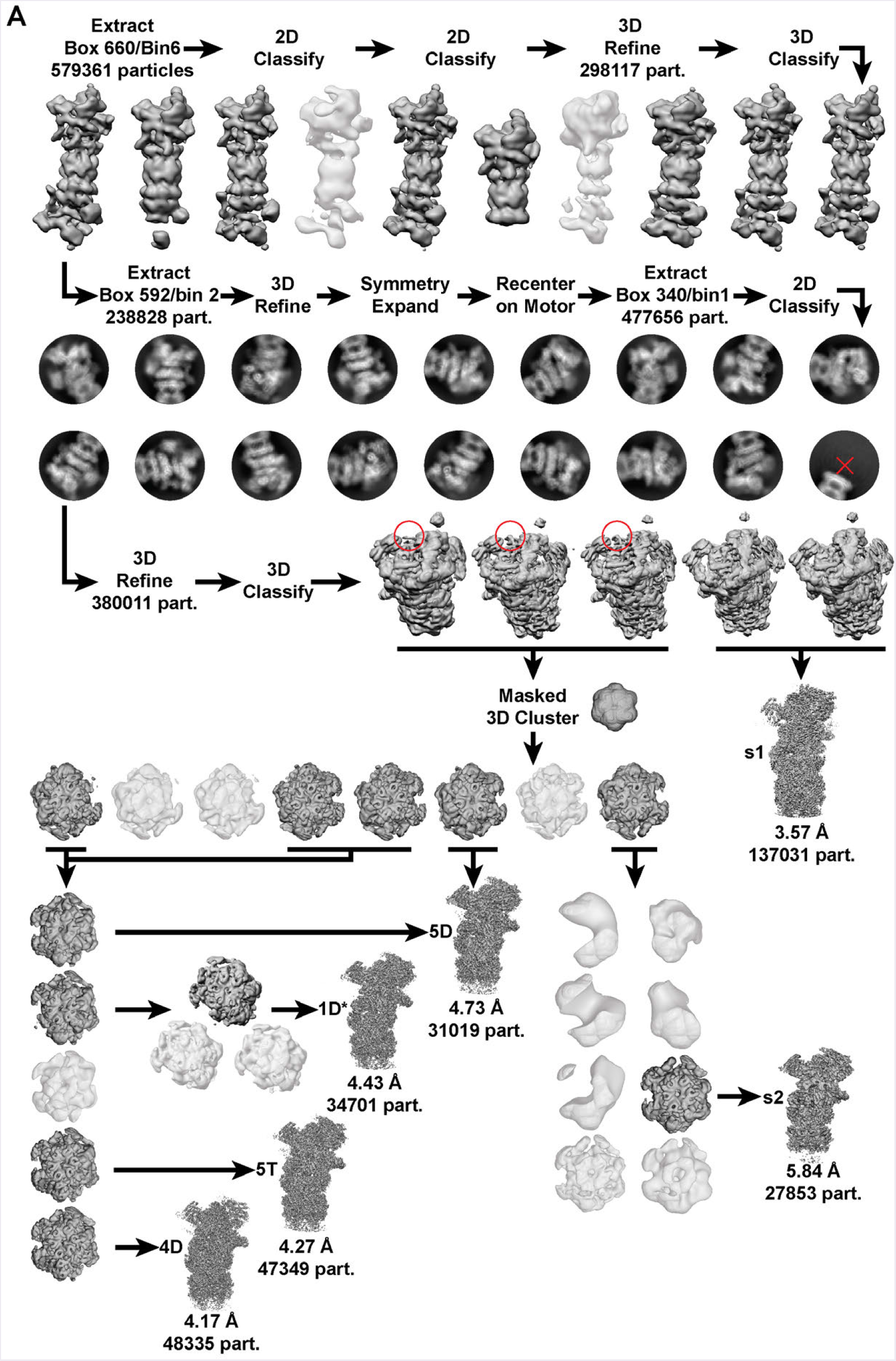
Schematic for cryo-EM single-particle data processing. Low resolution or artefactual reconstructions (white) and corresponding particles were excluded from subsequent processing steps, whereas all other reconstructions (gray) and corresponding particles were utilized. Representative 2D classes (dark circular background) are shown, and excluded 2D classes are denoted by a red cross. Globular ubiquitin-like density is indicated by a red circle.

**Supplementary Figure 2B.**
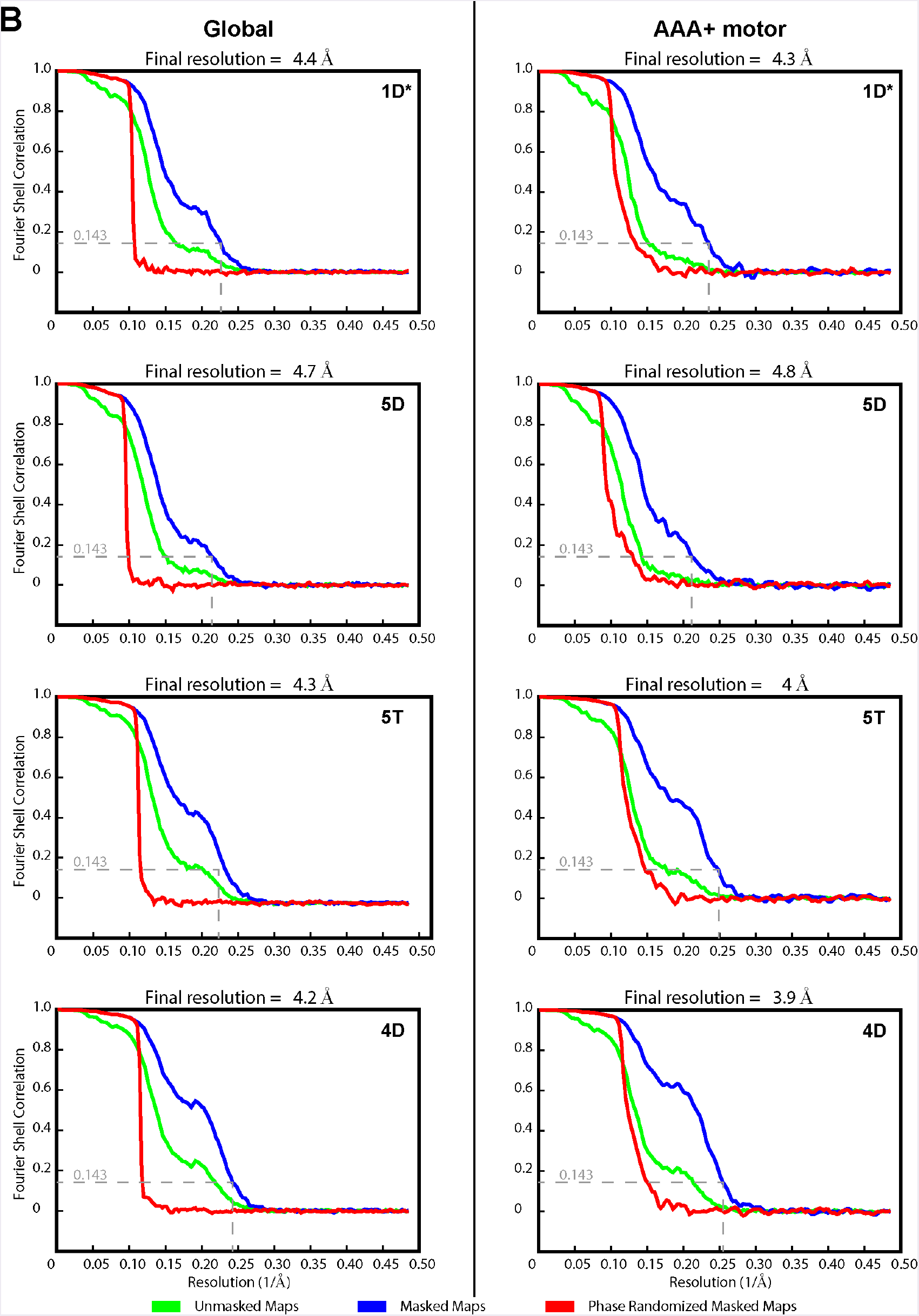
Gold Standard FSC for the 1D*, 5D, 5T, and 4D reconstructions. The Fourier Shell Correlation (FSC) curves are shown for the unmasked (green), masked (blue), and phase randomized (red) reconstructions. The left column corresponds to the global reconstructions, the right column to the local (AAA+ motor focus refinement) reconstructions. The resolution at 0.143 FSC is indicated by a dashed line.

**Supplementary Figure 2C.**
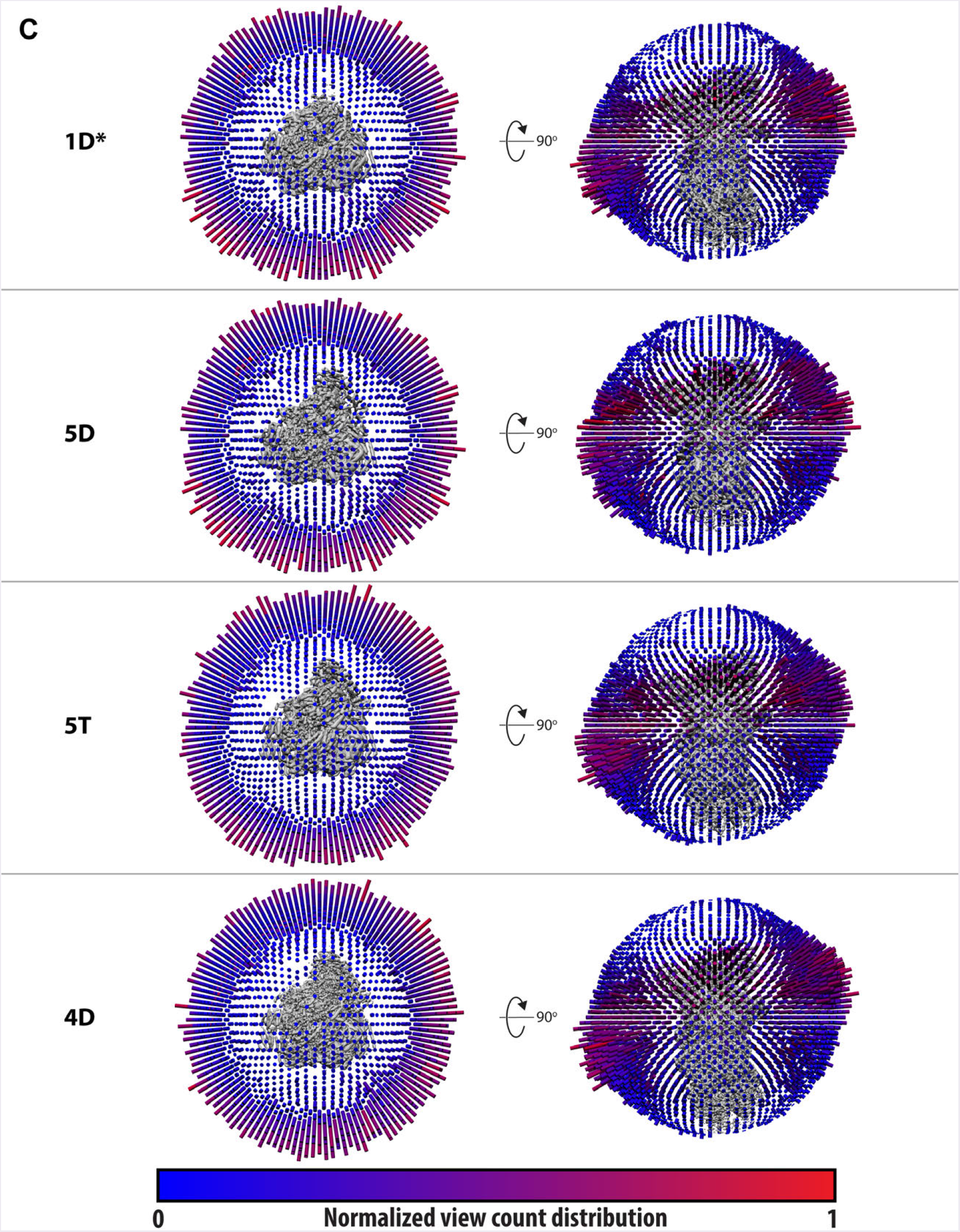
Distribution of Euler angles for the 1D*, 5D, 5T, and 4D reconstructions. Orthogonal views for each of the reconstructions are shown. The angular distributions are shown as columns whose longitudinal axis aligns with the normal of the corresponding back-projection (generated by RELION)^56,57^.

**Supplementary Figure 2D.**
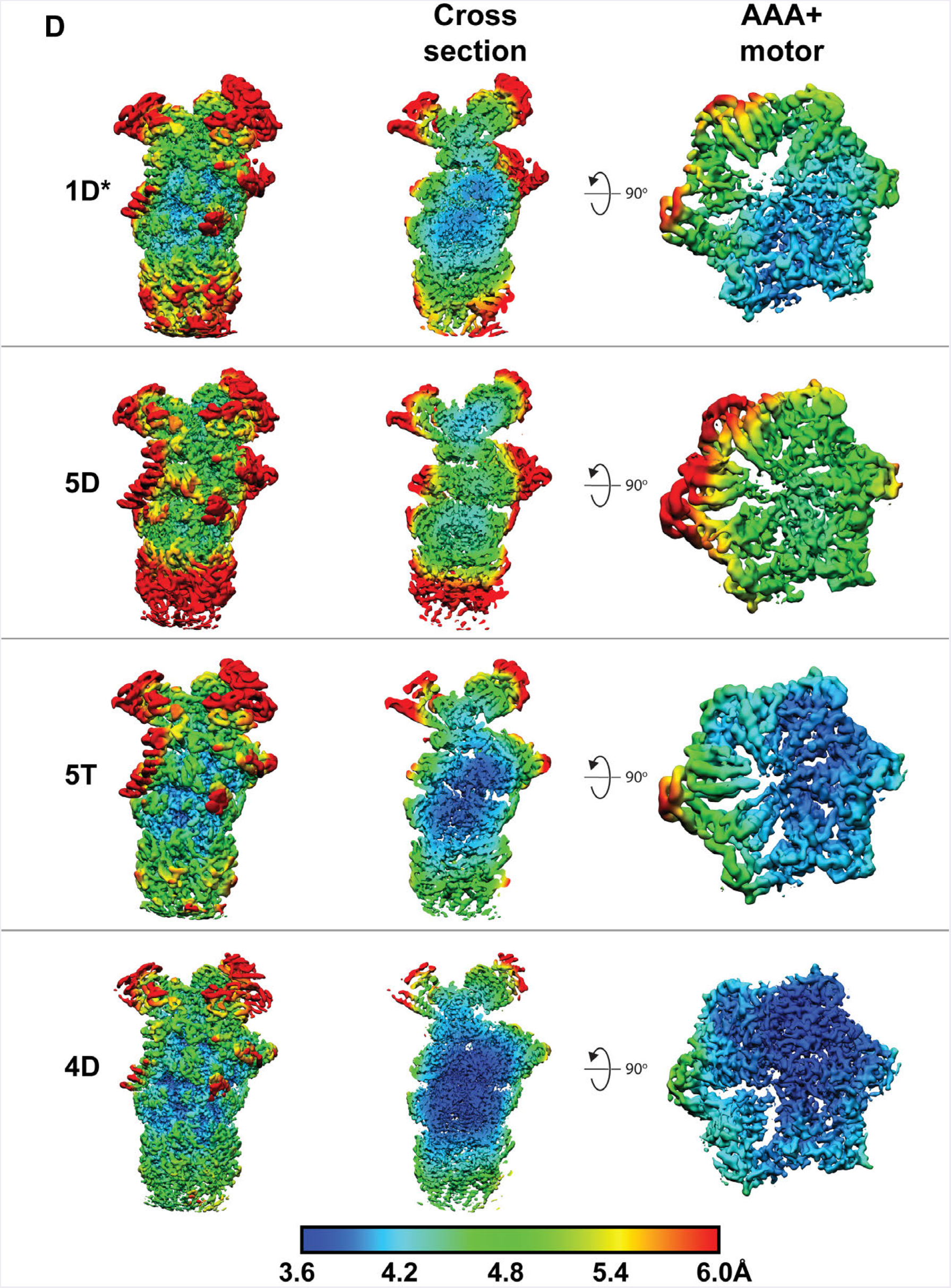
Local resolution estimates for the 1D*, 5D, 5T, 4D reconstructions. Local resolution estimates of the global reconstructions were calculated with RELION, and shown on the complete volume (left), on a coronal cross-section (middle), and on the AAA+ motor (right)^56,57^.

**Supplementary Figure 2E.**
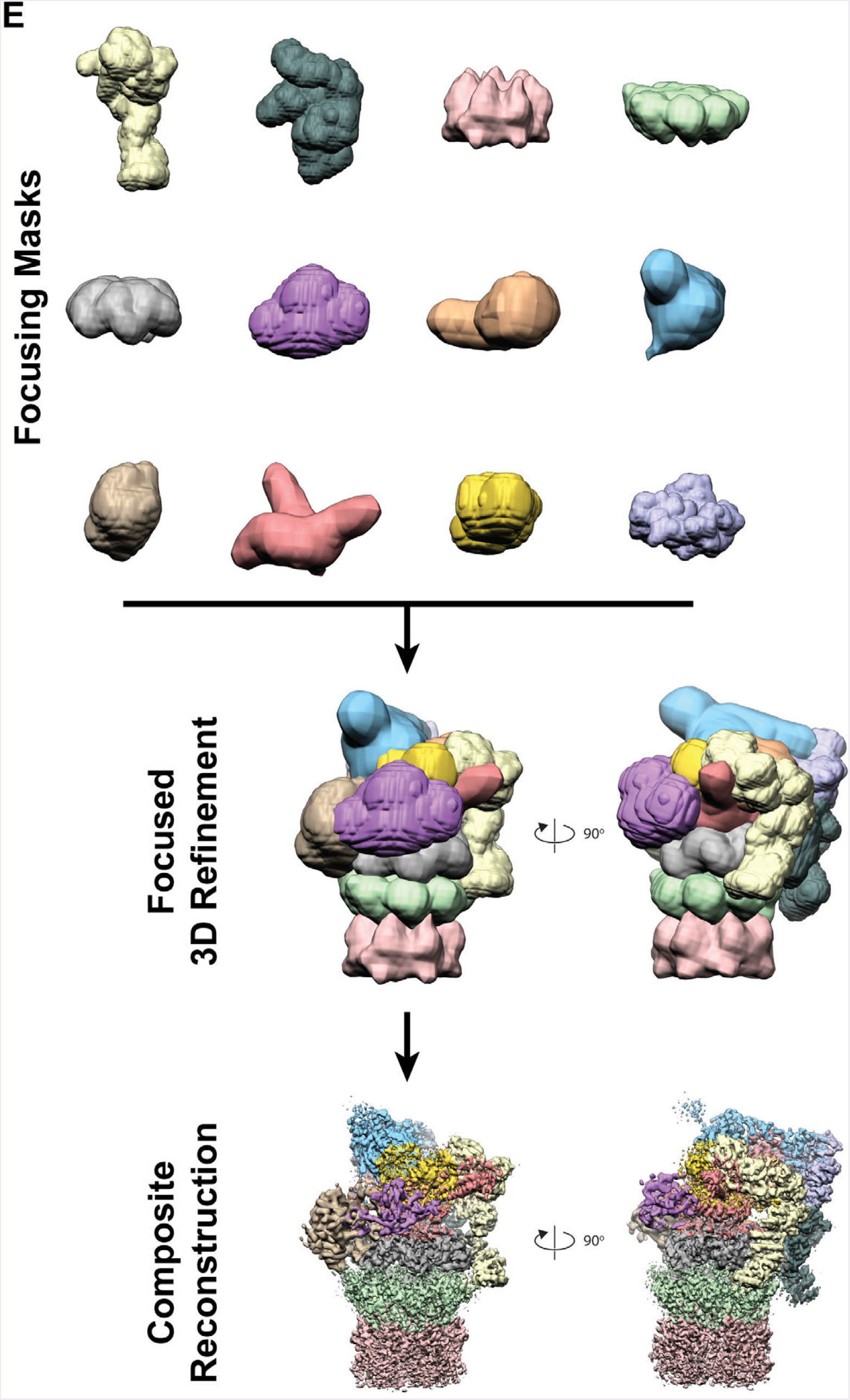
Procedure utilized to create 12-subvolume composite reconstructions for each 1D*, 5D, 5T, and 4D states. Twelve overlapping and inclusive masks with complete proteasomal coverage (top). The masks are utilized for focused 3D refinement (middle). Following 3D refinement, the reconstructions are re-masked, stitched together using the “ vop max” operator in UCSF Chimera^63^, and padded to the original global reconstruction dimensions. Overlay of the focused maps prior to stitching are shown at the bottom.

**Supplementary Figure 2F.**
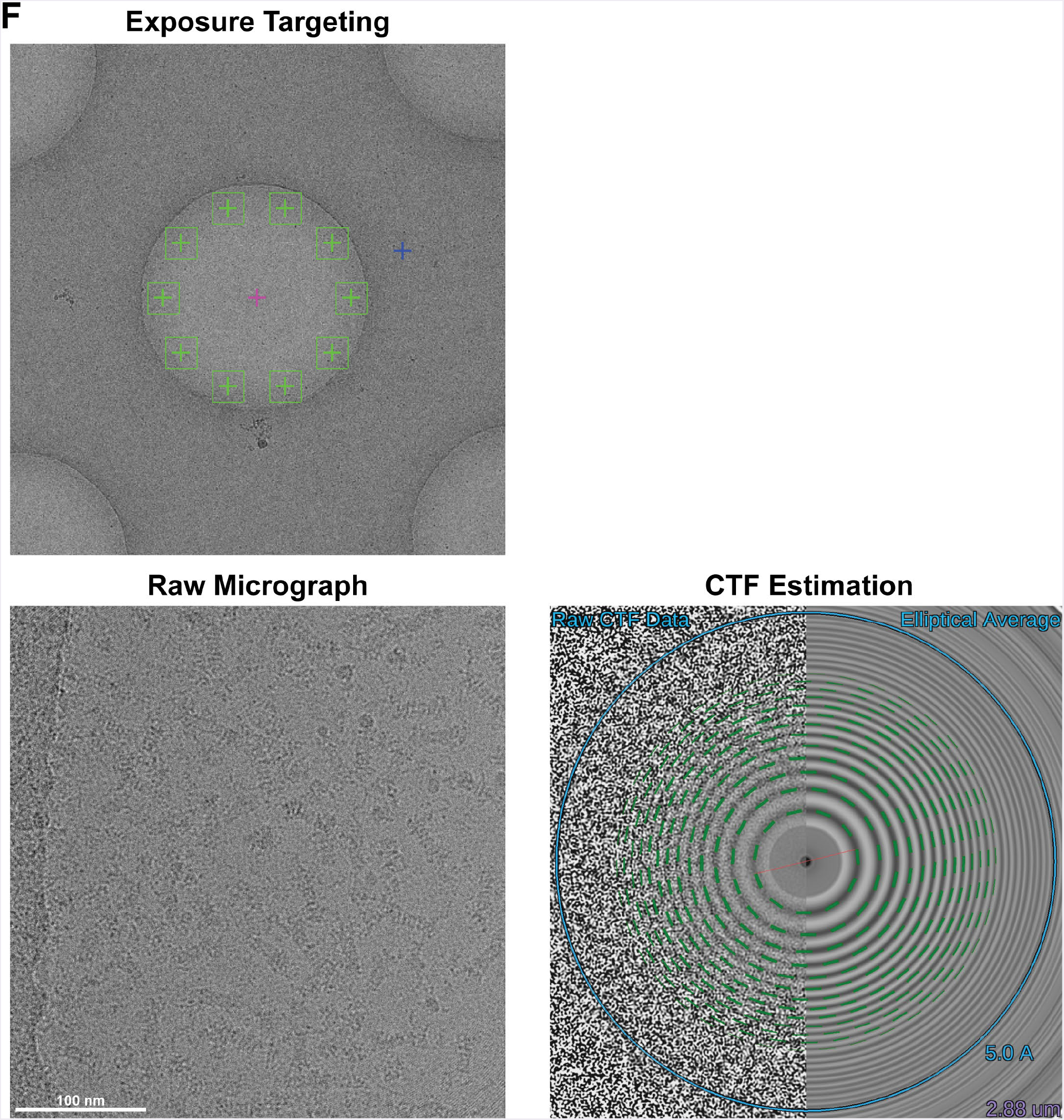
Cryo-EM Data collection. Ten micrographs (green squares, top left) were collected per focus routine (blue cross), after navigating to the center of a hole (pink cross). A representative micrograph (bottom left) and its corresponding CTF estimate are shown (bottom right). The micrograph depicts particle localization at the periphery of 2 μm holes.

**Supplementary Figure 2G.**
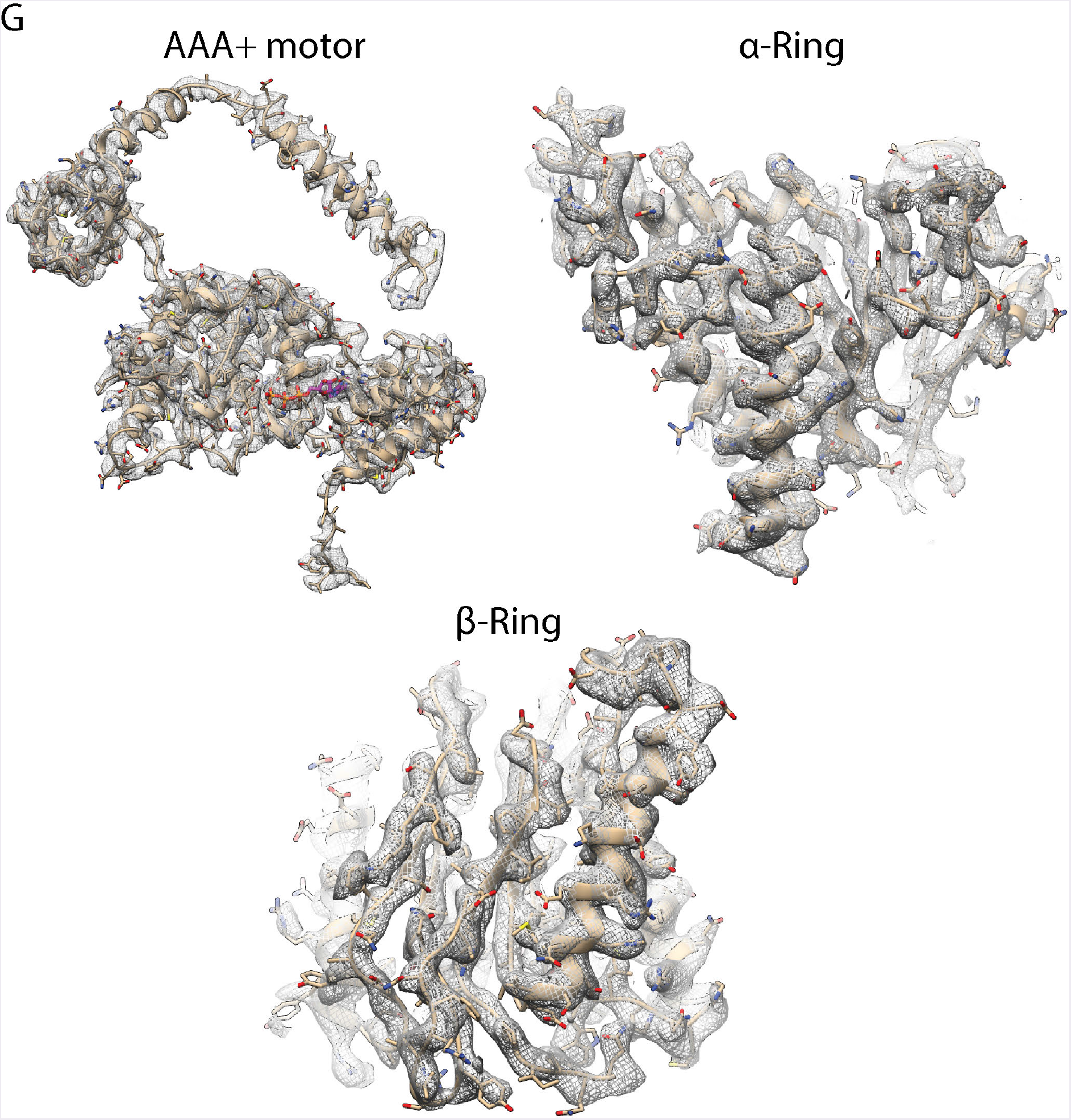
Representative high-resolution cryo-EM density. Density (gray mesh) for three distinct regions of the 4D state are shown with a corresponding atomic model (tan).

**Supplementary Figure 3.**
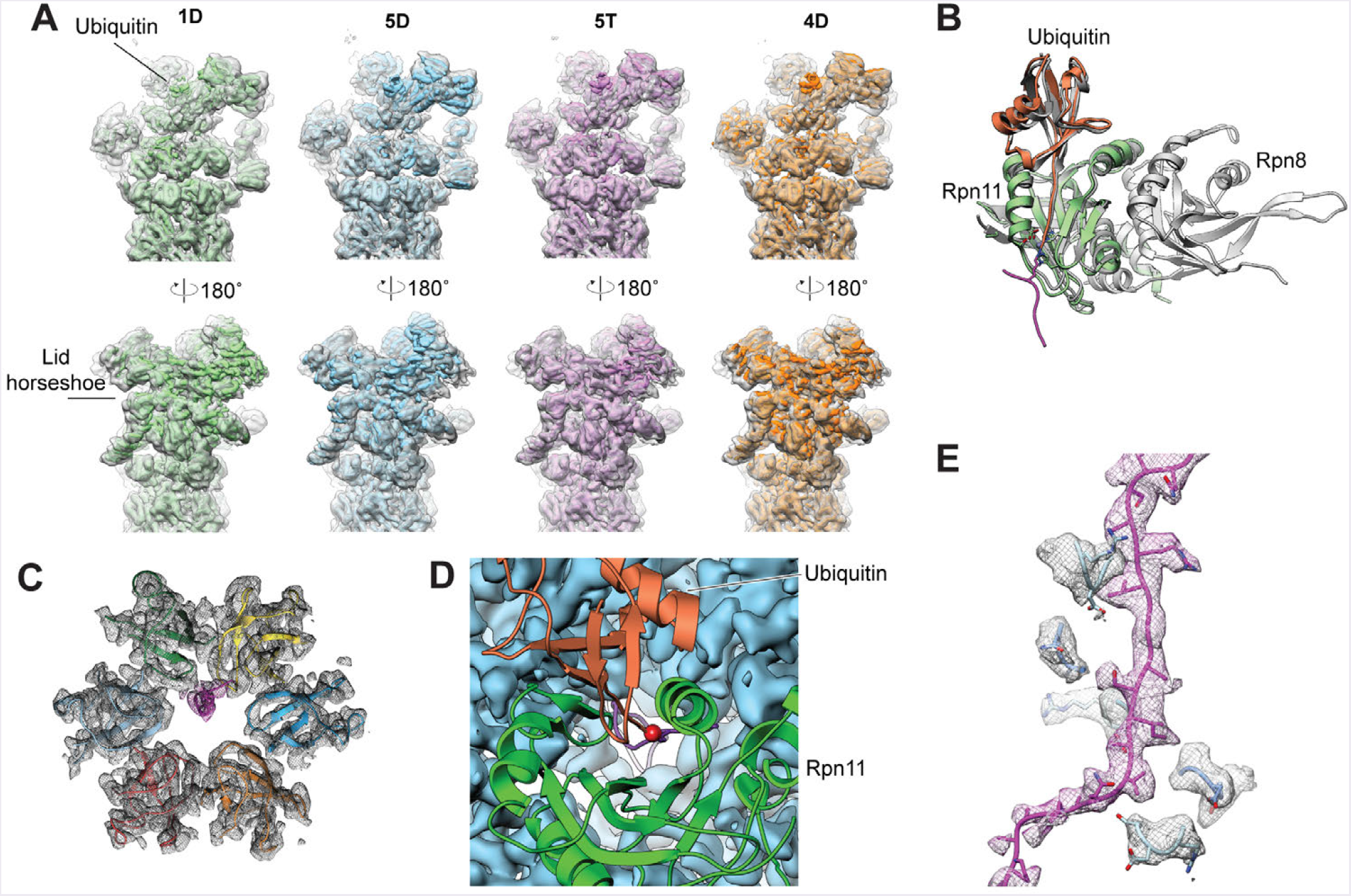
Stalled 26S proteasomes make extensive contacts with the stalled substrate. (A) Cryo-EM density for the 1D*, 5D, 5T and 4D states with their core particles aligned to the substrate-free s4 conformation (EMDB: 35 37^33^). (B) Alignment of the substrate-engaged 4D state (Rpn11 in green, ubiquitin in orange, and substrate in magenta) with the crystal structure of the ubiquitin-bound Rpn11/Rpn8 dimer (gray, PDB: 5U4P^38^). (C) Axial view of the N-ring demonstrating asymmetric placement of the substrate in its central pore. The substrate (magenta) is spatially closer to the OB folds of Rpt2 (yellow) and Rpt1 (green) than to the OB folds of Rpt6 (blue), Rpt3 (orange), Rpt4 (red), and Rpt5 (light blue). (D) Top view of the 4D state, looking through the central pore from above Rpn11 (green ribbon). Ubiquitin (orange ribbon) sits in the catalytic groove of Rpn11 directly above the pore of the motor (blue density). The isopeptide linkage (red sphere) aligns longitudinally with the central pore and the substrate’ s (magenta ribbon) translocation trajectory through the AAA+ motor. (E) Spiral staircase of pore-2 loops (blue) surrounding the substrate polypeptide (magenta) in the 4D state, with Cryo-EM density shown in mesh representation.

**Supplementary Figure 4A.**
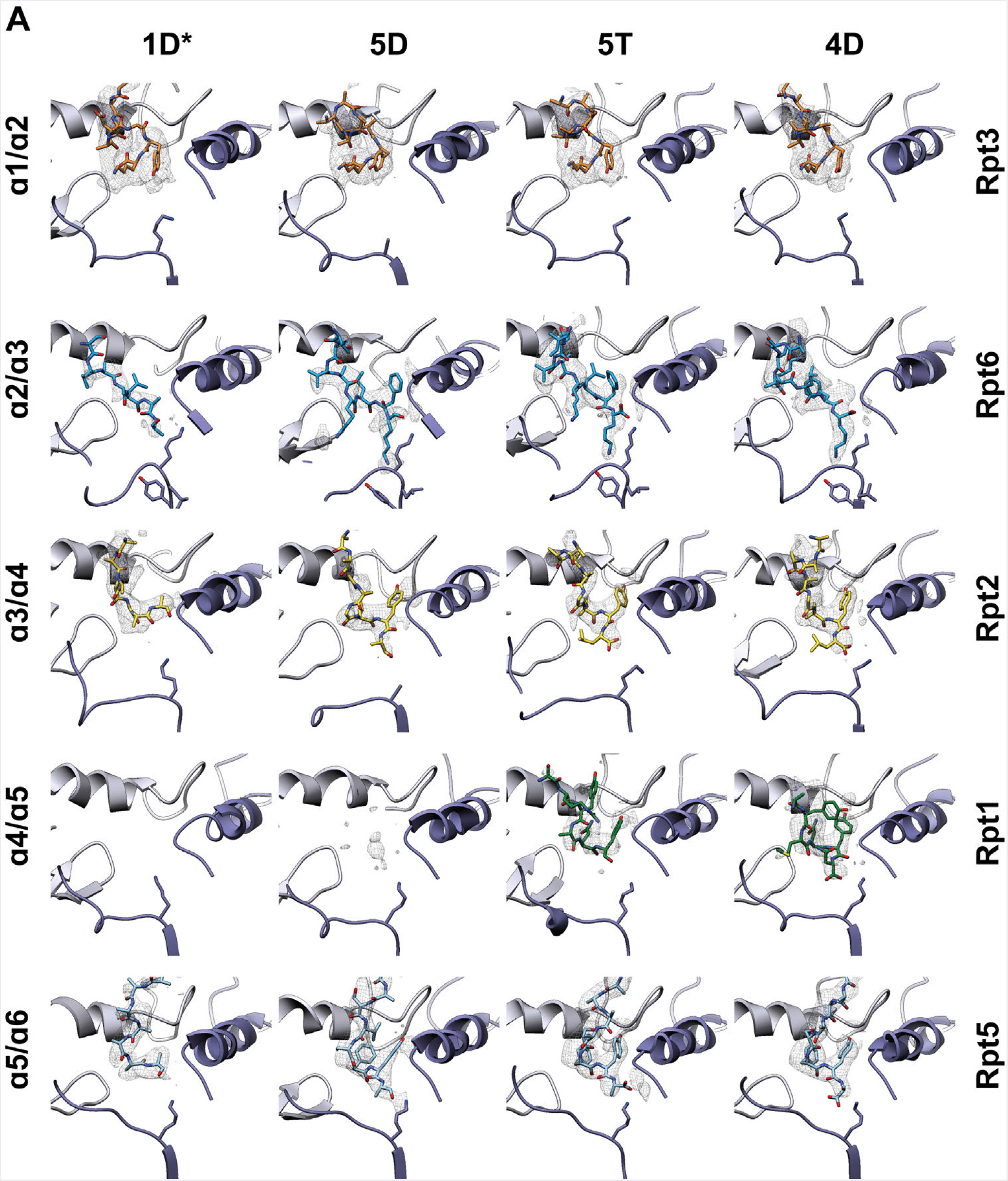
20S CP gate opening triggered by docking of Rpt C-terminal tails. The C-terminal tails of Rpt3 (orange), Rpt6 (blue), Rpt2 (yellow), Rpt1 (green), and Rpt5 (light blue) are shown in the pockets formed by neighboring CP α-subunits (α1-α6). Columns correspond to the four substrate-engaged states in the order of the motor progressing through the ATPase cycle.

**Supplementary Figure 4B-D.**
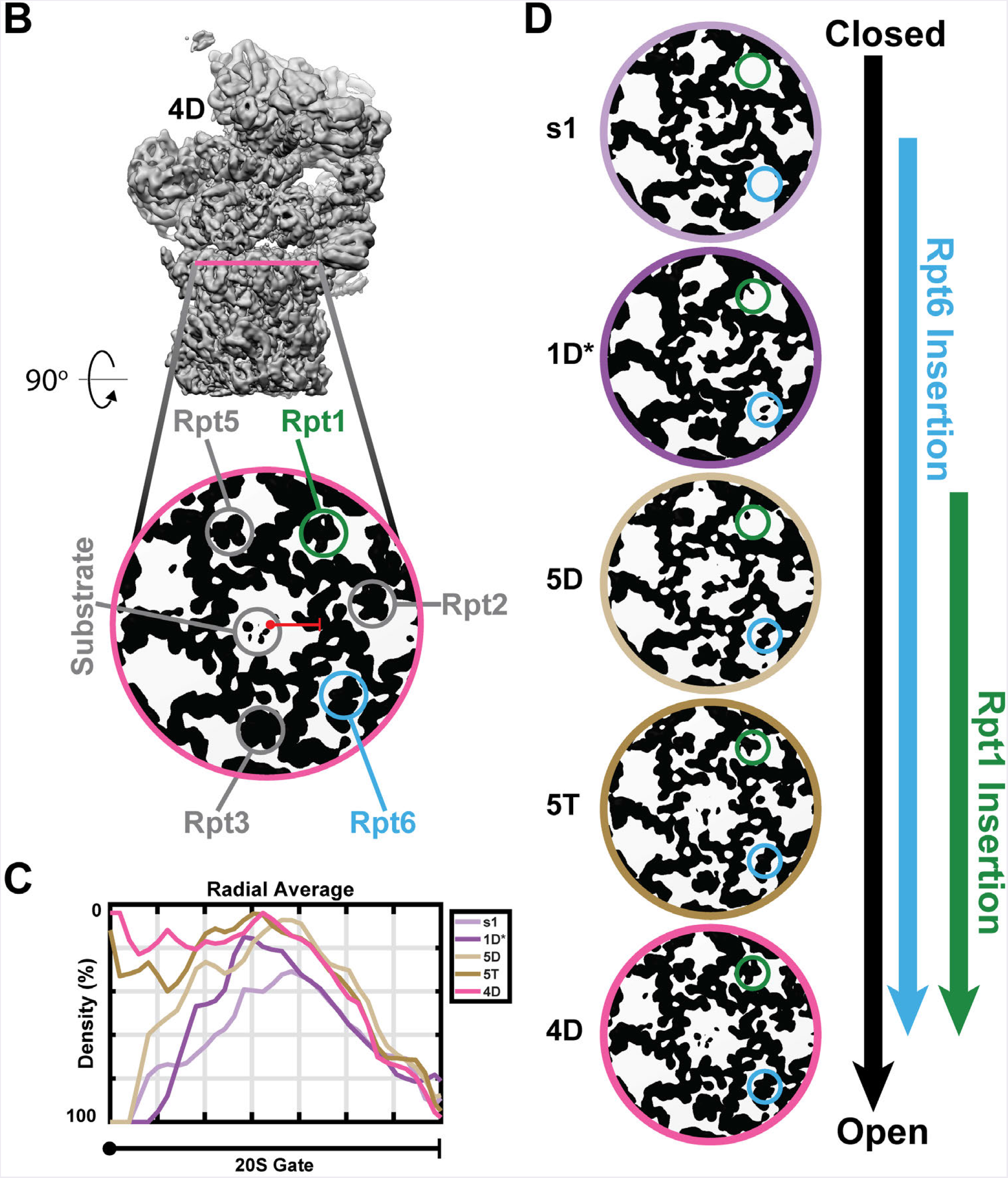
Differential CP-gate opening conformations. (B) The plane of the CP gate is indicated by a pink line in the side view of the proteasome in the 4D state. Shown below is the corresponding slice of the EM density, shown as a binary representation with CP gate density as black over a white background. The invariably docked C-terminal tails of Rpt2, Rpt3, and Rpt5 circled in gray, and the density corresponding to the C-terminal tails of Rpt1 and Rpt6 circled in green and blue, respectively. The portion used to calculate the radial average plotted in (C) is designated with a red line. (C) Radial average of a binary representation of CP gate density for the published s1 state (EMDB: 3534,^33^ and our four substrate-engaged states. (D) CP gate densities for each proteasome state, plotted from the most closed to the most open gate. Density changes for the docked tails of Rpt1 and Rpt6 are denoted with green and blue circles, respectively.

**Supplementary Figure 4E.**
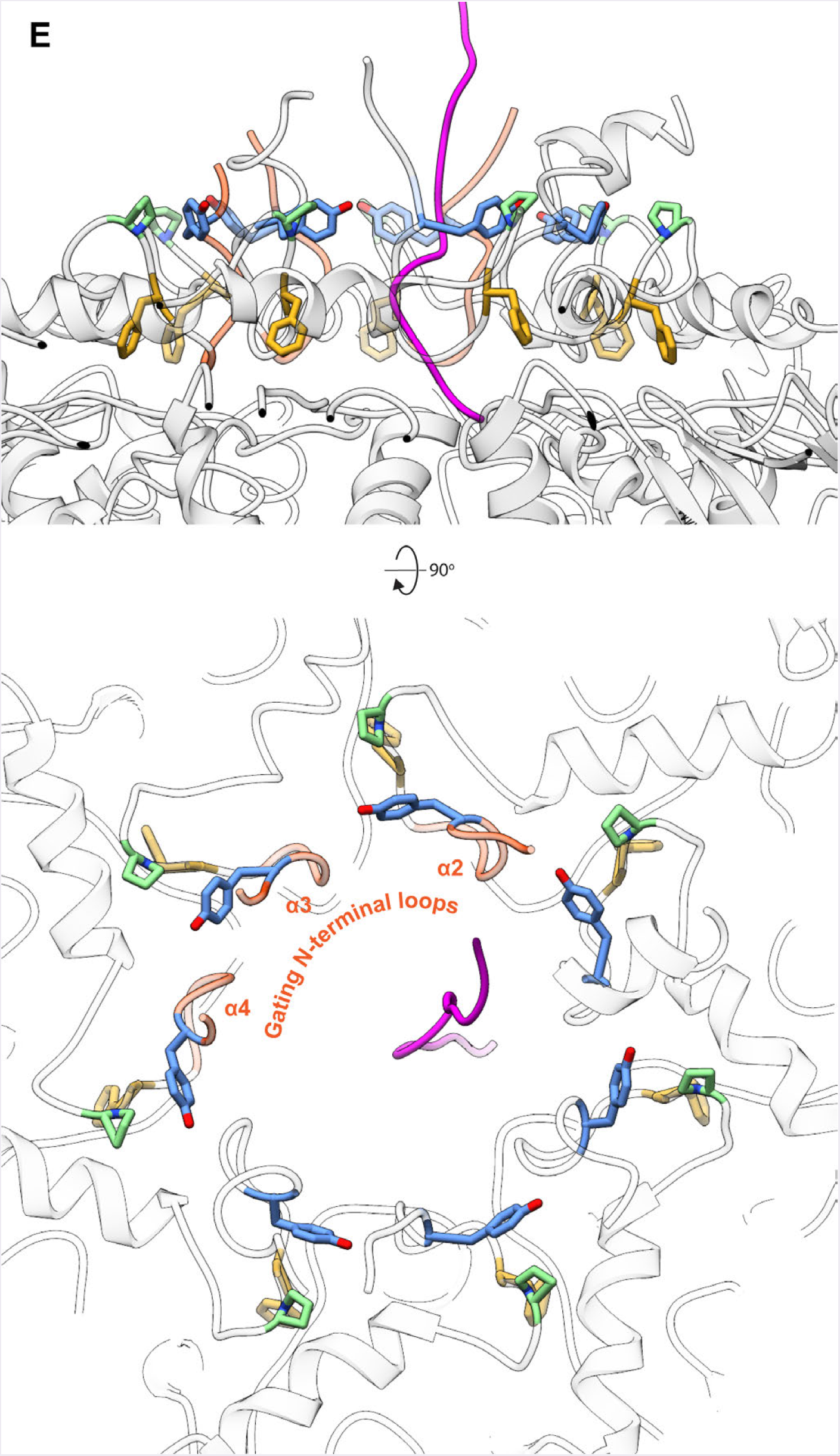
Fully open-gate conformation in the 4D state. The substrate polypeptide extends past the AAA+ motor and into the open CP gate. A hydrophobic collar formed by Tyr residues in the N-terminal tails of α-subunits is surrounding the CP access pore. A conserved Pro residue (green) is positioned within CH/π interaction distance from each Tyr residue, while a Phe residue (tan) neighboring the Pro points towards the peptidase core.

**Supplementary Figure 5A,B.**
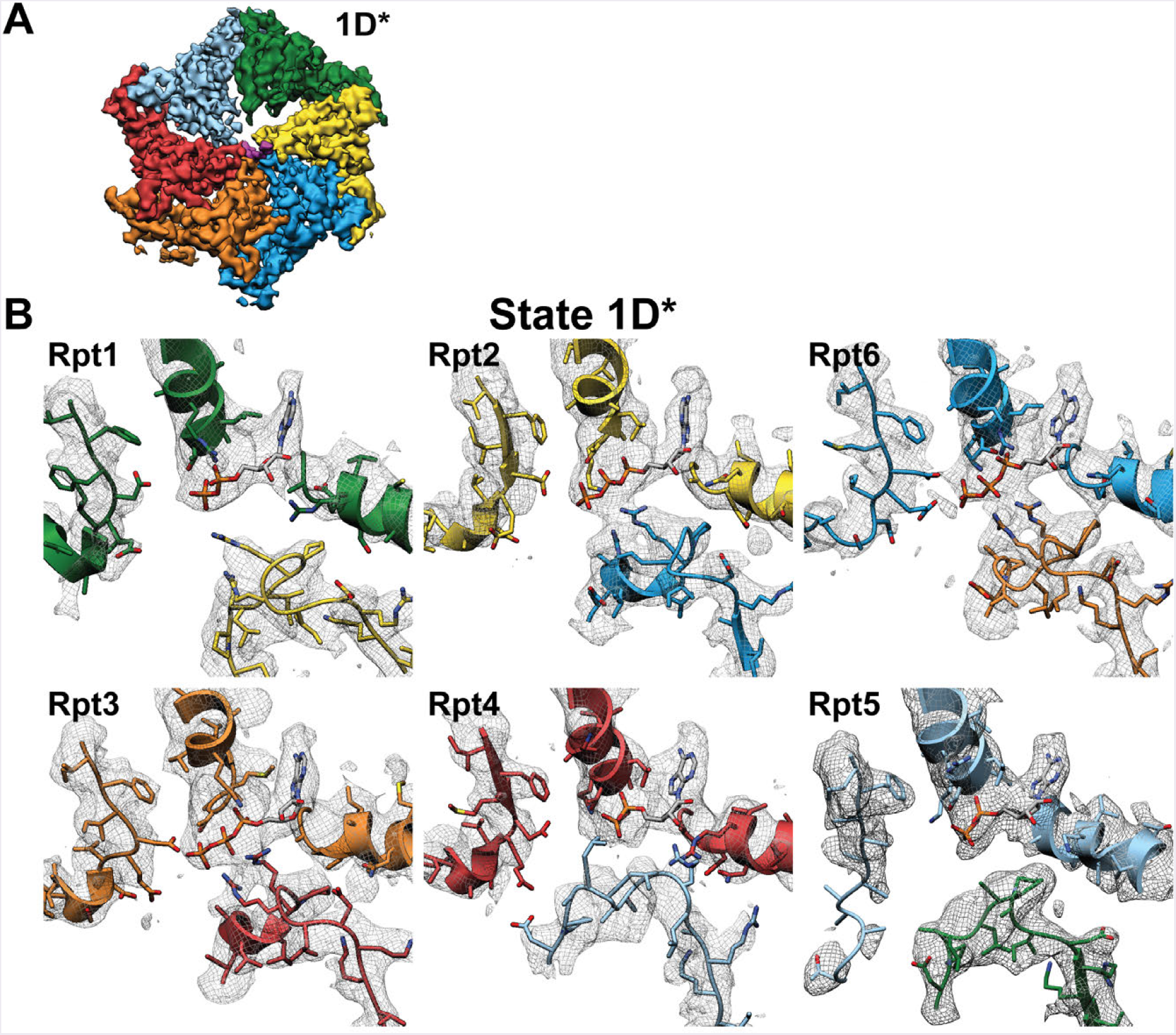
Nucleotide identities in binding pockets of the 1D* state. (A) Top view of AAA+-motor density in the 1D* state. Substrate density (magenta) is shown in the central pore formed by AAA+ motor subunits Rpt1 (green), Rpt2 (yellow), Rpt6 (blue), Rpt3 (orange), Rpt4 (red), and Rpt5 (light blue). (B) Individual nucleotide-binding pockets shown for each Rpt subunit and its clockwise-next neighbor in the 1D* state (colored by Rpt), with corresponding density in gray mesh.

**Supplementary Figure 5C,D.**
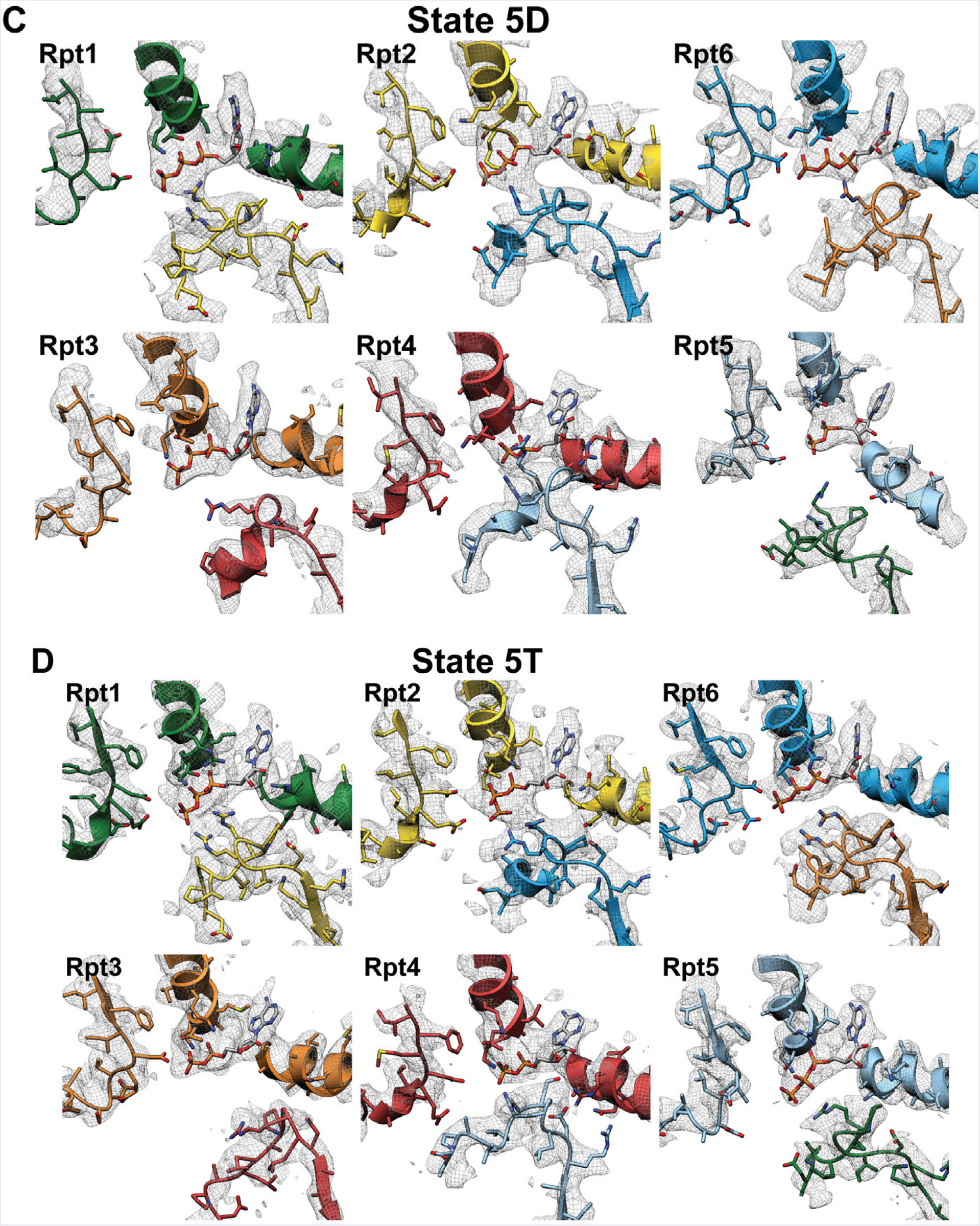
5C,D. Nucleotide identity in the binding pockets of the 5D and 5T states. (C) Individual nucleotide-binding pockets for each Rpt and its clockwise-next neighbor in the 5D state, colored by Rpt and with the corresponding density shown as gray mesh. (D) Equivalent binding pockets in the 5T state.

**SupplementaryFigure 5E,F.**
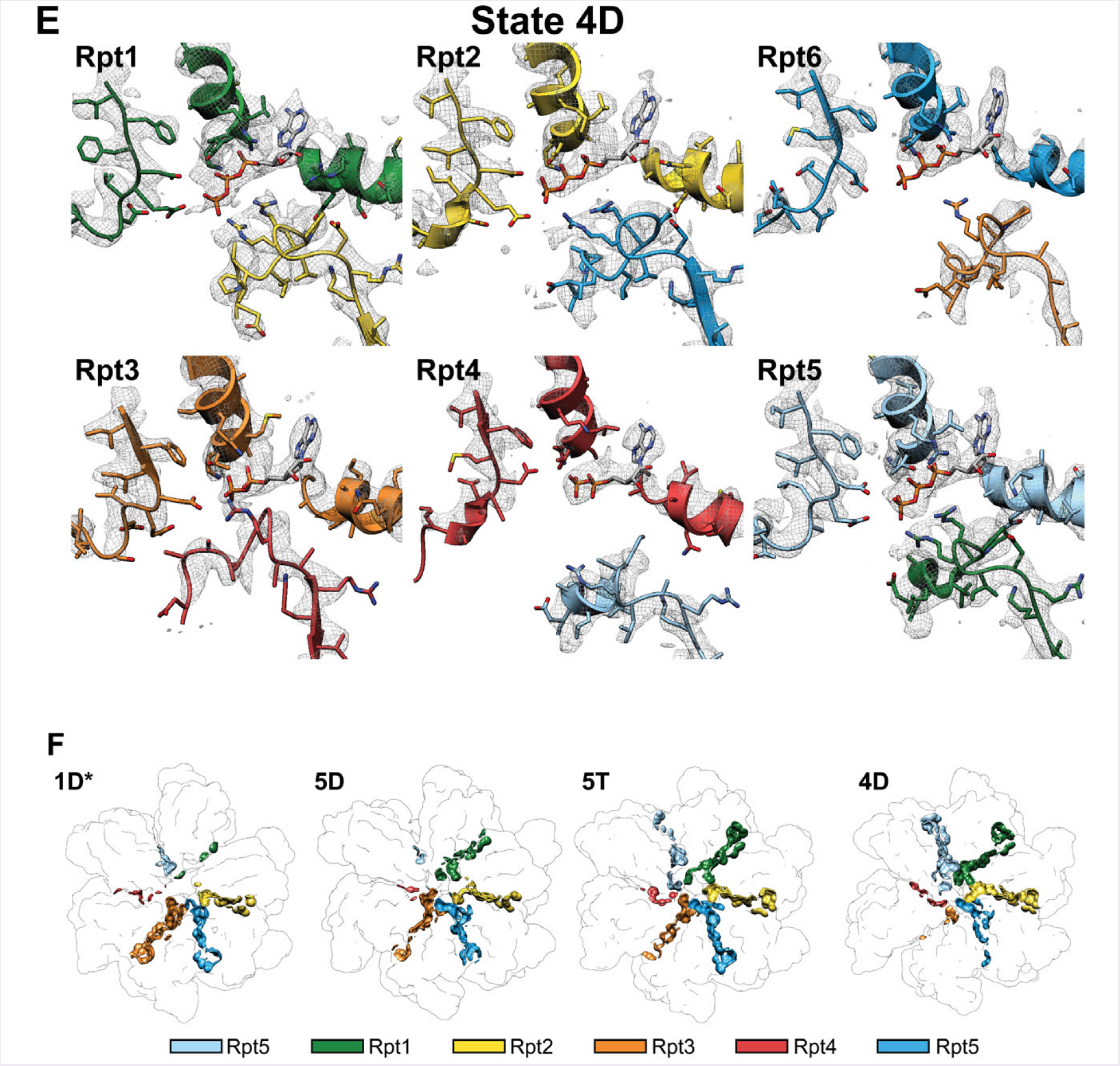
Nucleotide identity in the nucleotide binding pockets of the 4D state. (E) Individual nucleotide-binding pockets for each Rpt and its clockwise-next neighbor in the 4D state, colored by Rpt and with the corresponding density shown as gray mesh. (F) Inter-subunit contact area (molecular surfaces that are within 2 A of each other) for each neighboring pair of large AAA+ subdomains. The contact area is colored according to the nucleotide-bound Rpt for each pair.

**SupplementaryFigure 6.**
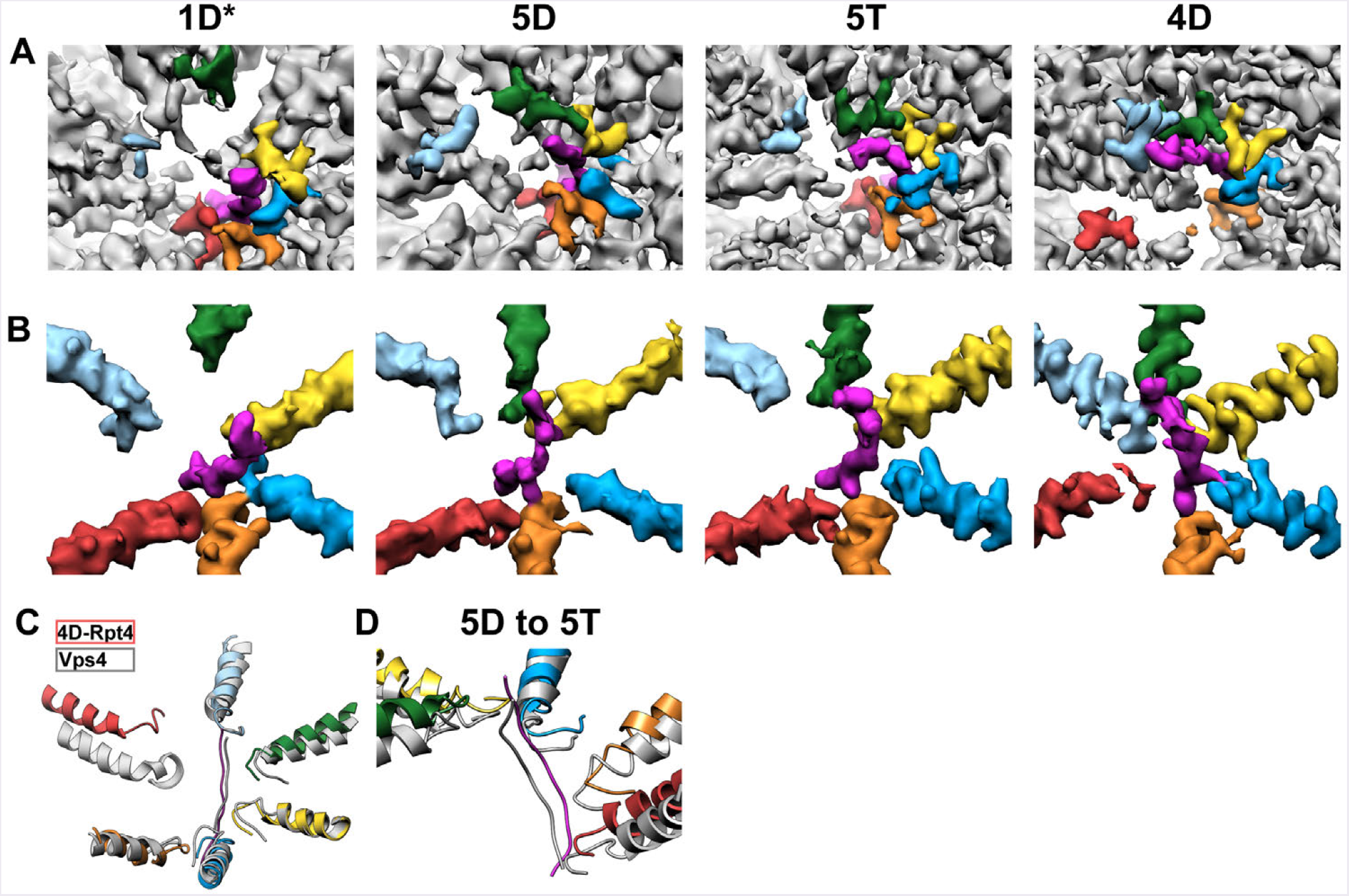
Spiral-staircase arrangements of pore-1 and pore-2 loops in contact with substrate. (A) Top view of the AAA+ motor for the 1D*, 5D, 5T and 4D states, with the pore-1 loop density colored by Rpt (Rpt1 in green, Rpt2 in yellow, Rpt6 in blue, Rpt3 in orange, Rpt4 in red, and Rpt6) and substrate density shown in magenta. (B) Densities for the pore-2 loop and helix 10 in the 1D*, 5D, 5T and 4D states, colored by Rpt. (C) Spiral staircase of pore-1 loops rotated and aligned for the 4D state and Vps4 (PDB: 6ap1^45^). (D) The 5D and 5T states, aligned by their core particles, show similar spiral staircase arrangements. Pore-1 loop and helix 8 are colored by Rpt for the 5D state and shown in gray for 5T.

**SupplementaryFigure 7.**
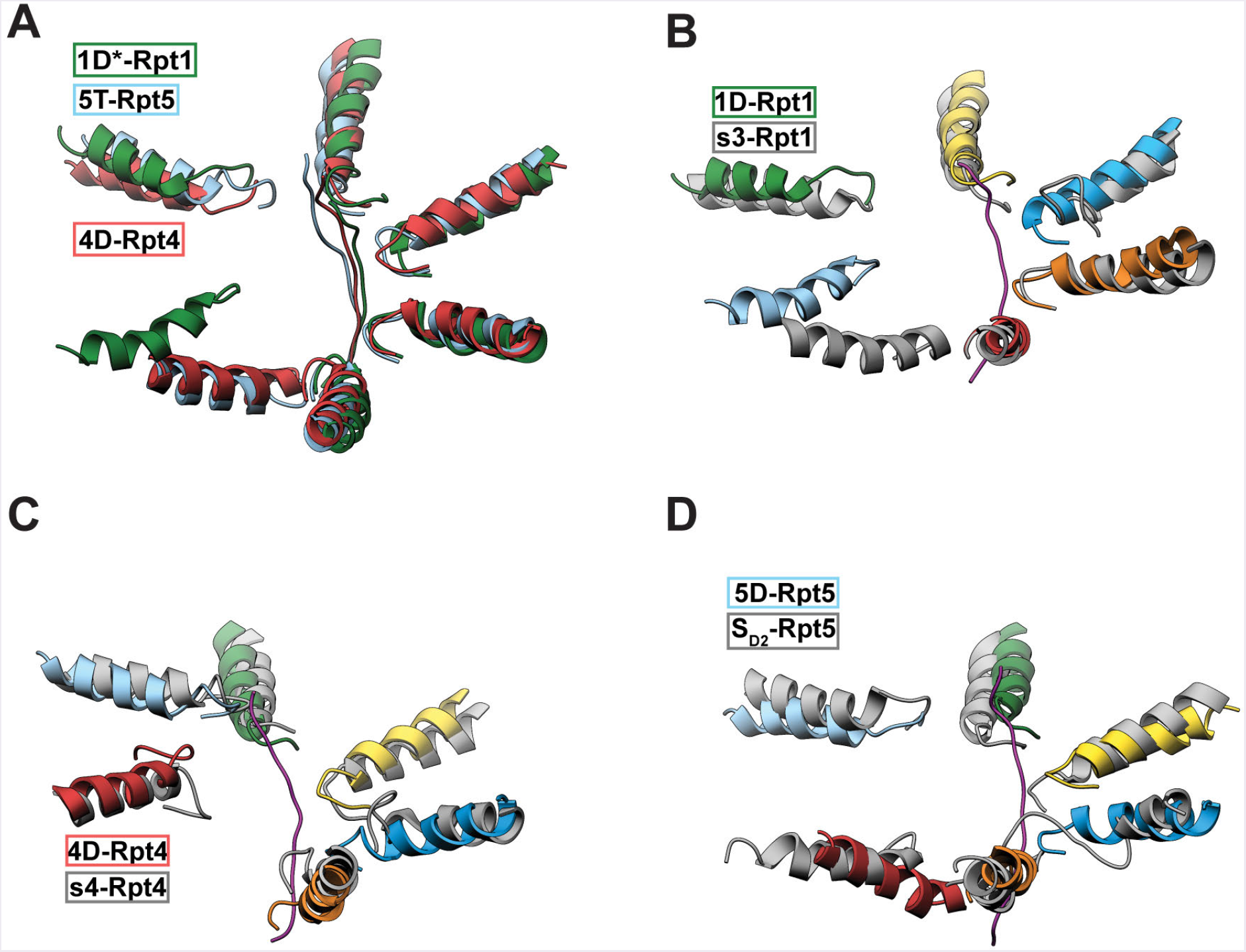
Overlays of spiral staircases from substrate-bound proteasome states with staircases from substrate-free proteasome structures. (A) Spiral staircases of the 1D* (dark green), 5T (light blue), and 4D (red) pore-1 loop helices rotated in increments of 60 degrees to align the spiral staircases, with a label indicating the overlaying disengaged subunits. (B) Rotated and aligned pore-1-loop spiral staircases for the 1D* and s3 (PDB:5mpb^33^) states, with 1D* colored by Rpt (Rpt1 in green, Rpt2 in yellow, Rpt6 in blue, Rpt3 in orange, Rpt4 in red, and Rpt6), substrate in magenta, and s3 in gray, and a label indicating the overlaying disengaged subunits. (C) Rotated and aligned spiral staircases of pore-1 loops for the 4D and s4 (PDB:5mpc^33^) states, with 4D colored by Rpt, substrate in magenta, and s4 in gray, and a label indicating the overlaying disengaged subunits. (D) Rotated and aligned spiral staircases of pore-1 loops for the 5D and SD2 (PDB:5vfq^35^) states, with 5D colored by Rpt, substrate in magenta, and SD2 in gray, and a label indicating the overlaying disengaged subunits.

**Supplementary Table 1.** Cryo-EM data collection, refinement, and validation statistics

| Name | State 1D* | State 5D | State 5T | State 4D |
| --- | --- | --- | --- | --- |
| PDB ID |  |  |  |  |
| EMDB ID |  |  |  |  |
| **Data collection and processing** |  |  |  |  |
| Microscope | Titan Krios | Titan Krios | Titan Krios | Titan Krios |
| Camera | K2 Summit | K2 Summit | K2 Summit | K2 Summit |
| Magnification | 29000 | 29000 | 29000 | 29000 |
| Voltage (kV) | 300 | 300 | 300 | 300 |
| Total electron fluence (e<sup>-</sup>/Å<sup>2</sup>) | 50 | 50 | 50 | 50 |
| Fluence rate (e<sup>-</sup>/pixel/sec) | 8 | 8 | 8 | 8 |
| Defocus range (μm) | -1.5 to -3.0 | -1.5 to -3.0 | -1.5 to -3.0 | -1.5 to -3.0 |
| Pixel size (Å/pixel)* | 1.03 | 1.03 | 1.03 | 1.03 |
| Micrographs collected (no.) | 11,664 | 11,664 | 11,664 | 11,664 |
| Total extracted picks (no.) | 579,361 | 579,361 | 579,361 | 579,361 |
| Refined particles (no.) | 298,117 | 298,117 | 298,117 | 298,117 |
| Final Particles (no.) | 34,701 | 31,019 | 47,349 | 48,335 |
| Symmetry | C1 | C1 | C1 | C1 |
| Global resolution (Å) |  |  |  |  |
| FSC 0.5 (unmasked/masked) | 7.9/6.7 | 8.4/7.2 | 7.5/5.9 | 7.3/4.3 |
| FSC 0.143 (unmasked/masked) | 5.5/4.4 | 6.7/4.7 | 5.0/4.3 | 4.6/4.2 |
| Local resolution range (Å) | 3.8-8.2 | 4.1-8.2 | 3.8-8.3 | 3.6-7.7 |
| **Model composition** |  |  |  |  |
| Nonhydrogen atoms | 24,767 | 23,659 | 24,663 | 44,272 |
| Protein residues | 3320 | 3294 | 3290 | 5824 |
| Ligands | 6 | 6 | 6 | 6 |
| **Refinement** |  |  |  |  |
| MapCC (global/local) | 0.52/0.782 | 0.46/0.80 | 0.521/0.80 | 0.66/0.80 |
| Map sharpening *B*-factor (Å<sup>2</sup>) | -146 | -147 | -134 | -140 |
| R.m.s. deviations |  |  |  |  |
| Bond lengths (Å) | 0.01 | 0.01 | 0.01 | 0.01 |
| Bond angles (°) | 0.98 | 0.91 | 1.01 | 1.02 |
| **Validation** |  |  |  |  |
| EMRinger<sup>64</sup> score | 1.67 | 1.28 | 2.55 | 2.96 |
| MolProbity score | 2.14 | 2.11 | 2.01 | 1.97 |
| Clashscore | 5.19 | 4.96 | 4.38 | 3.69 |
| Poor rotamers (%) | 0 | 0 | 0 | 0 |
| Ramachandran plot |  |  |  |  |
| Favored (%) | 83.07 | 84.03 | 85.40 | 85.22 |
| Allowed (%) | 16.90 | 15.94 | 14.60 | 14.74 |
| Disallowed (%) | 0.03 | 0.03 | 0.00 | 0.03 |

\* Calibrated pixel size at the detector

**Supplementary Table 2.** Pocket openness and staircase measurements for Figures 2 and 3

|  | **Rpt1** | **Rpt2** | **Rpt6** | **Rpt3** | **Rpt4** | **Rpt5** |
| --- | --- | --- | --- | --- | --- | --- |
| Distance from Walker A to ISS Helix (Å) |  |  |  |  |  |  |
| **1D\*** | 21.9 | 12.2 | 15.0 | 15.3 | 20.5 | 21.5 |
| **5D** | 16.7 | 12.6 | 14.5 | 15.7 | 21.1 | 24.0 |
| **5T** | 14.8 | 11.1 | 14.5 | 16.0 | 22.4 | 14.6 |
| **4D** | 14.6 | 10.2 | 14.1 | 23.4 | 22.1 | 13.0 |
| Distance from Walker A to Arg Finger (Å) |  |  |  |  |  |  |
| **1D\*** | 16.6 | 13.8 | 14.1 | 12.5 | 14.8 | 16.5 |
| **5D** | 14.1 | 12.9 | 12.2 | 13.1 | 15.2 | 21.9 |
| **5T** | 13.6 | 13.7 | 14.0 | 17.0 | 16.9 | 13.8 |
| **4D** | 13.8 | 13.0 | 14.4 | 17.1 | 18.2 | 13.3 |
| Contact Area between Large Domains of Rpt Subunits (Å<sup>2</sup>) |  |  |  |  |  |  |
| **1D\*** | 70.1 | 410.6 | 513.9 | 648.3 | 71.8 | 156.4 |
| **5D** | 347.4 | 452.4 | 494.2 | 398.8 | 43.8 | 66.4 |
| **5T** | 608.4 | 606.7 | 682.8 | 373.7 | 93.8 | 317.7 |
| **4D** | 784.3 | 699.3 | 343.0 | 74.3 | 121.2 | 697.2 |
| Distance from Pore-1 loop to Core Plane* (Å) |  |  |  |  |  |  |
| **1D\*** | 52.4 | 52.8 | 48.1 | 42.8 | 37.9 | 48.4 |
| **5D** | 51.8 | 48.1 | 41.9 | 36.9 | 31.3 | 50.0 |
| **5T** | 50.3 | 46.6 | 41.6 | 36.2 | 28.9 | 51.0 |
| **4D** | 46.8 | 42.4 | 36.7 | 30.6 | 49.3 | 52.3 |

\*The plane defined by the $\alpha$ helices encircling the gate of the core residues at $\alpha 1$ 16-28, $\alpha 2$ 18-30, $\alpha 3$ 18-30, $\alpha 4$ 15-27, $\alpha 5$ 13-25, $\alpha 6$ 18-30, $\alpha 7$ 17-29

**Movie 1: Tracing the substrate path in the engaged 26S proteasome structure**. The high-resolution cryo-EM structure of the substrate-engaged 26S proteasome (state 4D shown) reveals the substrate-attached ubiquitin moiety bound to Rpn11 and shows the substrate polypeptide traversing the AAA+ motor, engaged by five pore-1 loops in a spiral-staircase arrangement and entering the open gate of the core peptidase.

**Movie 2: Transition between the consecutive 5D, 5T and 4D states of the substrate-engaged 26S proteasome**. A conformational morph between the 5D, 5T, and 4D motor states reveals nucleotide exchange, ATP hydrolysis, and substrate translocation steps. Substrate is colored magenta and the Rpts are colored by subunit, with Rpt1 in green, Rpt2 in yellow, Rpt6 in blue, Rpt3 in orange, Rpt4 in red, and Rpt5 in light blue.

